# Single Cell Track and Trace: live cell labelling and temporal transcriptomics via nanobiopsy

**DOI:** 10.1101/2023.06.13.544323

**Authors:** Fabio Marcuccio, Chalmers C. Chau, Georgette Tanner, Marilena Elpidorou, Martina A. Finetti, Shoaib Ajaib, Morag Taylor, Carolina Lascelles, Ian Carr, Iain Macaulay, Lucy F. Stead, Paolo Actis

**Author notes:** Joint corresponding authors. Correspondence to and. These authors contributed equally (Fabio Marcuccio, Chalmers C. Chau, Georgette Tanner).

## Abstract

Single-cell RNA sequencing has revolutionised our understanding of cellular heterogeneity, but whether using isolated cells or more recent spatial transcriptomics approaches, these methods require isolation and lysis of the cell under investigation. This provides a snapshot of the cell transcriptome from which dynamic trajectories, such as those that trigger cell state transitions, can only be inferred. Here, we present cellular nanobiopsy: a platform that enables simultaneous labelling and sampling from a single cell without killing it. The technique is based on scanning ion conductance microscopy (SICM) and uses a double-barrel nanopipette to inject a fluorescent dye and to extract femtolitre-volumes of cytosol. We used the nanobiopsy to longitudinally profile the transcriptome of single glioblastoma (GBM) brain tumour cells in vitro over 72hrs with and without standard treatment. Our results suggest that treatment either induces or selects for more transcriptionally stable cells. We envision the nanobiopsy will transform standard single-cell transcriptomics from a static analysis into a dynamic and temporal assay.

## INTRODUCTION

Microfluidics platforms coupled with advances in molecular biology have enabled high-throughput analysis of the minute amount of genetic material contained in a single cell^1^. The advent of spatial transcriptomics has then further enabled the generation of single cell gene expression data while maintaining the spatial context of those cells within a tissue^2^. However, all these approaches are end-point assays that require the cell to be lysed or fixed before profiling. An exciting new frontier in single cell analysis is the facilitation of temporal/sequential/longitudinal cellular profiling ^3,4^. Chen, Guillaume-Gentil et al established Live-seq, a technique based on fluidic force microscopy^5^ allowing the pressure-driven RNA extraction from living cells at distinct time points^6^. The authors sequentially profiled the transcriptomes of individual macrophages before and after lipopolysaccharide (LPS) stimulation, and of adipose stromal cells pre- and post- differentiation^6^. Nadappuram, Cadinu et al developed minimally invasive dielectrophoretic nanotweezers based on dual carbon nanoelectrodes that allowed the trapping and extraction of nucleic acids and organelles from living cells without affecting their viability^7^. The authors demonstrated the single organelles resolution of their techniques but did not demonstrate longitudinal sampling. Alternative approaches relying on vertical nanoneedle platforms have the potential to enable high throughput single cell manipulation^8^ and they recently enabled the intracellular sampling from a few cells to investigate cellular heterogeneity ^9^. Our group has pioneered the use of nanopipettes coupled with electrowetting and integrated into a Scanning Ion Conductance Microscope (SICM)^10,11^ to perform nanobiopsies of living cells in culture^12^. SICM relies on the measurement of the ion current between an electrode inserted in a glass nanopipette, the probe, and a reference electrode immersed in an electrolytic solution where the cells are placed^11^. By applying a voltage between the two electrodes, an ion current flows through the nanopore at the tip of the nanopipette. When the nanopipette approaches a surface, the measured ion current drops. This current drop is proportional to the separation between the nanopore and the sample and can be used as active feedback to maintain the nanopipette-sample distance constant and it has been extensively used for high resolution topographical mapping of living cells ^11^. The nanobiopsy technique has enabled the extraction of RNA and organelles from single cells, the study of mRNA compartmentalization within neuronal cells^13^ and the localised sampling of mitochondria from human tissues^14^.

Here, we show the development of a platform technology enabling nanoinjection of exogenous molecules into a living cell and extraction of femtolitre-volumes of cytoplasm, via a double-barrel nanopipette. The platform enables longitudinal nanobiopsy of the same cell to profile gene expression changes over a 72-hour period. As a proof of concept, we applied our novel method to investigate changes in gene expression in a model of the most aggressive brain cancer, glioblastoma (GBM) caused by non-surgical elements of the standardised treatment given to patients: radiotherapy and chemotherapy with the drug temozolomide (TMZ). This approach allowed for direct characterisation of therapy driven changes in cell state, offering unique insights into the gene expression changes that may underpin the ability of these cells to phenotypically adapt and resist treatment. Such investigations allow for unprecedented insights into how cells respond under different conditions or with different treatments, which will be invaluable in many research settings.

## RESULTS

Double-barrel nanopipettes integrated into a SICM enable the injection into living cells followed by extraction of minute amount of the cytoplasm for downstream gene expression profiling. Our platform technology employs a double-barrel nanopipette (individual barrel with a pore of ∼150 nm, **Supplementary Fig. SF1.1**) where one barrel, filled with an organic solution is used as an electrochemical syringe to perform cytoplasmic extractions ^12–16^, while the second barrel, filled with an aqueous electrolyte solution, provides a stable ion current for precise positioning and enables the nanoinjection of exogenous molecules into the cell. **Figure 1** illustrates the nanobiopsy technology and its workflow composed of automated nanopipette approach to the cell (a), penetration of the cell membrane followed by nanoinjection (b), extraction of cytoplasmic RNA via nanobiopsy (c), transcriptomics analysis based on next-generation sequencing and bioinformatics analysis (d).

**Figure 1.**
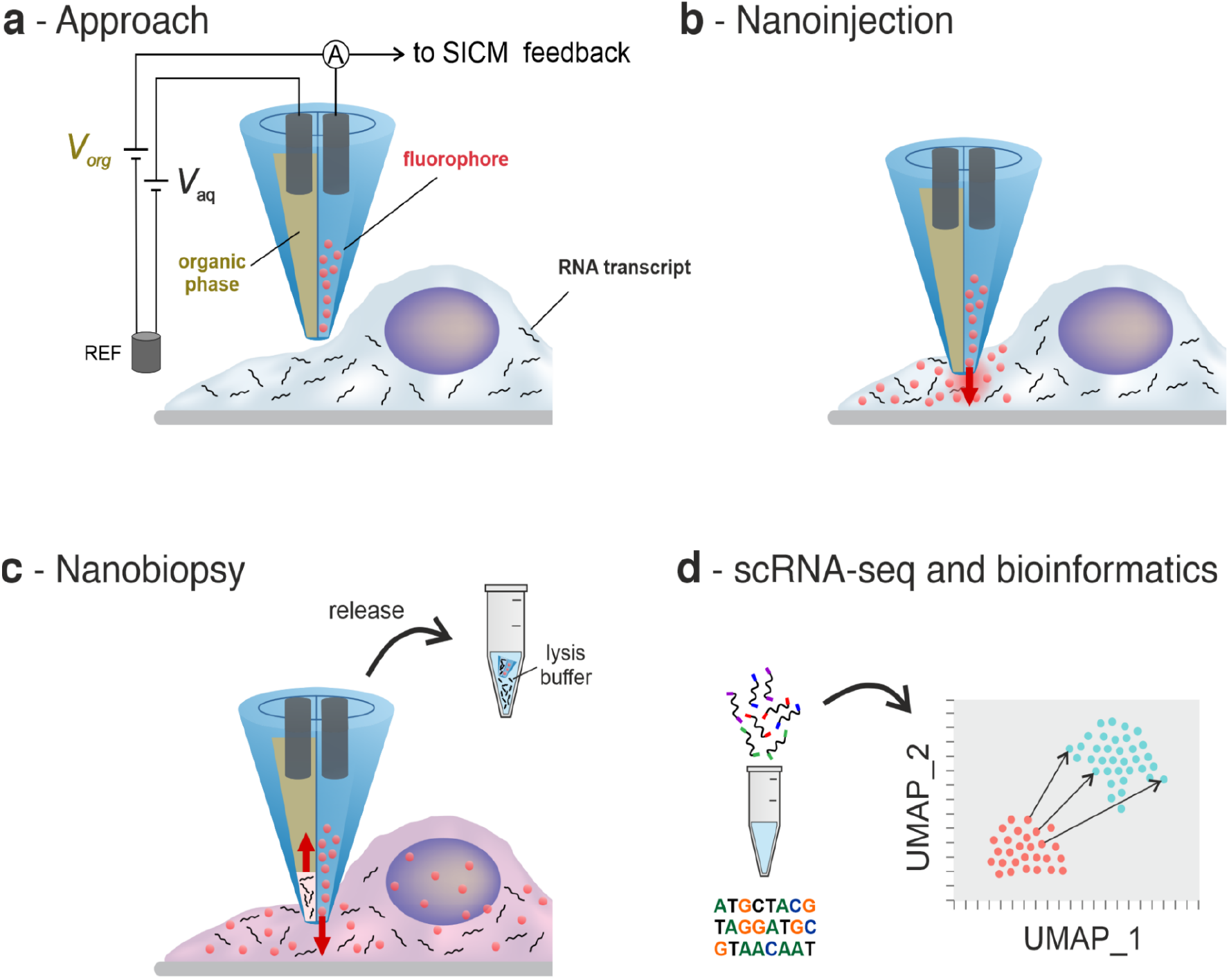
Illustration of the main phases of the single-cell nanobiopsy procedure enabling cytoplasmic nanoinjections and extractions from individual cells. A double-barrel nanopipette integrated to an SICM is brought in proximity of the cellular membrane (a) where it penetrates the latter enabling the release of the fluorophore through the aqueous barrel (b), and cytoplasmic extractions through the organic barrel (c). After the extraction, the sample containing RNA transcripts is released in lysis buffer and it is reverse transcribed, amplified and sequenced for downstream gene expression analysis via bioinformatics (d).

### APPROACH

In the approach phase, the double-barrel nanopipette automatically approaches the cell membrane under control of the SICM interface^17^. The electrolyte solution in the aqueous barrel is characterised by an electrical conductivity κ_aq_ = 1.30 S/m which is two orders of magnitude higher than the conductivity of the organic solution κ_org_ = 0.01 S/m^15^. This results in a median electrical resistance of the aqueous barrel R_aq_ = 45.7 MΩ being 59 times smaller than the median resistance of the organic barrel R_org_ = 2.7 GΩ which translates with higher values of ion current magnitude that facilitate the SICM operations. A positive bias is applied to the electrode in the aqueous barrel (*V*_*aq*_ *=* 200 mV) with respect to a reference electrode immersed in the culture medium. When the nanopipette approaches the plasma membrane, the magnitude of the ion current decreases due to the increase in the nanopipette access resistance^10,18^ until the setpoint current (99.5% of the baseline current magnitude) is detected.

At this point, the SICM feedback mechanism stops and retracts the nanopipette to keep a 2 μm distance from the cell membrane. The ion current through the aqueous barrel *i*_aq_ corresponding to the vertical position *z* of the nanopipette recorded during the approach phase, the boxplots showing the resistance of the organic and aqueous barrel, and the optical micrographs of the process are shown in **Supplementary Fig. SF1.2 and SF1.3**. For the entire duration of the approach phase, the potential applied to the electrode in the organic phase is kept at a constant positive value (*V*_org_ = 300 mV) to prevent the aqueous phase from entering the nanopipette ^12,15^.

### NANOINJECTION

In the nanoinjection phase, the nanopipette position is manually controlled to enable the reproducible penetration of the cell membrane. First the nanopipette is lowered by 2 μm to reach the cell membrane, then the potential applied to the aqueous barrel is switched to *V*_aq_ = -500 mV to allow the electrophoretic release of the anionic fluorophore from the nanopipette. Next, the nanopipette is lowered at high speed (100 nm/ms) with 100-nm steps until we detect a ∼2% drop in the ion-current magnitude that we assign to the nanopipette touching the cell membrane (subphase 2: membrane touch). The nanopipette is then further lowered until we detect a vibrational noise, which could be due to the proximity of the nanopipette tip to the petri dish (subphase 3: vibrational noise). A further 100-nm step brings the nanopipette in contact with the polymer coverslip of the petri dish and no ion current is detected (*i*_aq_ = 0 pA). Finally, the nanopipette is retracted by 100 nm and it is kept at a fixed position for approximately 1 minute to allow the release of a sufficient amount of fluorophore into the cytoplasm. This method is robust and reproducible, allowing penetration of the membranes of different cell types (HeLa: epithelial cell; M049K: glial cell) with distinct mechanical properties^19,20^, to enable injection of exogenous molecules and extraction of cytoplasmic content. **Figure 2a** shows the ion current *i*_aq_ and the potential applied *V*_aq_ in the aqueous barrel and the vertical position *z* of the nanopipette during nanoinjection, while the zoomed-in traces for the individual approach (1), membrane touch (2) and vibrational noise (3) phases are shown in **Fig. 2b**. Note that the nanopipette vertical position *z* increases as the distance between nanopipette and cell membrane decreases.

**Figure 2.**
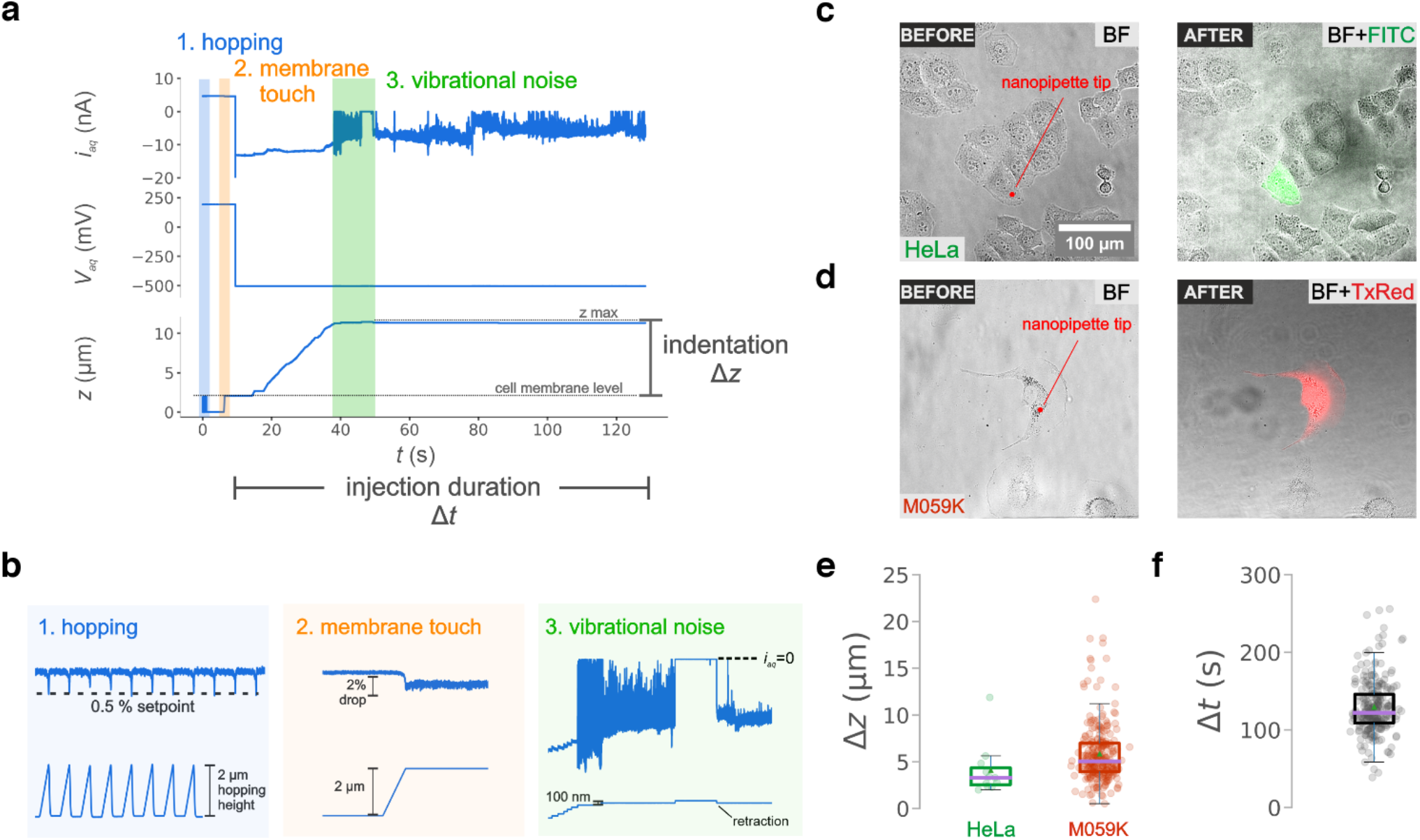
(a) Electrical traces for the ion current *i*_aq_ and potential bias *V*_aq_ in the aqueous barrel and z-position of the nanopipette *z* during the nanoinjection phase where the shaded regions highlight the hopping (1), membrane touch (2) and vibrational noise (3) subphases. The parameters for the indentation Δ*z* and the injection duration Δ*t* can be extracted. (b) Zoomed-in traces for the ion current *i*_aq_ (top) and nanopipette vertical position *z* (bottom) recorded during the hopping (1), membrane touch (2) and vibrational noise (3) subphases whose detection guarantees a successful membrane penetration and nanoinjection. The bright field (BF) and epifluorescence (FITC, TxRed) micrographs of a HeLa (c) and M059K (d) cell acquired before and after the nanoinjection phase show the cell emitting a fluorescent signal following injection of a green (ATTO 488) and red (ATTO 565) fluorophore, respectively. (e) Boxplots showing the maximum indentation Δ*z* in the case of nanoinjections performed into HeLa (n=10) and M059K cells (n=256), and (f) the total injection duration Δ*t* (n=256).

Also, the power spectral density of the ion current *i*_aq_ traces can be used as quality control because when the presence of the individual sub-phases strongly correlated with a successful nanoinjection (**Supplementary Section S1.4-S1.5, Supplementary Figs. SF1.4 - SF1.6**) resulting into a fluorescently labelled cell. **Figures 2c-d** show the optical micrographs obtained before and after the nanoinjection of a green anionic fluorophore (ATTO 488) into a HeLa cell and a red anionic fluorophore (ATTO 565) into a M059K cell. **Figure 2e** shows the indentation of the nanopipette onto the cellular membrane necessary to puncture the membrane and access the cytoplasm extracted from the traces shown in **Fig. 2a**. The boxplots show a median value (purple line) of Δ*z* = 3.32 μm and Δ*z* = 5.06 μm and an average (green triangle) of Δ*z* = 4.12 μm and Δ*z* = 4.85 μm in the case of HeLa and M059K cells, respectively. The distribution for total injection duration Δ*t* is shown in the boxplot in **Fig. 2f** where the median value is Δ*t* = 129 s.

### NANOBIOPSY

Following the nanoinjection step, our platform technology enables the extraction of intracellular RNA. **Figure 3a** shows an example of the ion current *i*_org_ and the potential *V*_org_ applied to the electrode in the organic barrel during the nanobiopsy phase where the electric potential is switched to *V*_org_ = -500 mV for 10 s. Upon switching the potential, the ion current in the organic barrel increases from *i*_org_ = 0.35 nA to *i*_org_ = - 2.83 nA and it decreases slowly until the end of the 10 s when the potential is returned to *V*_org_ = 300 mV.

**Figure 3.**
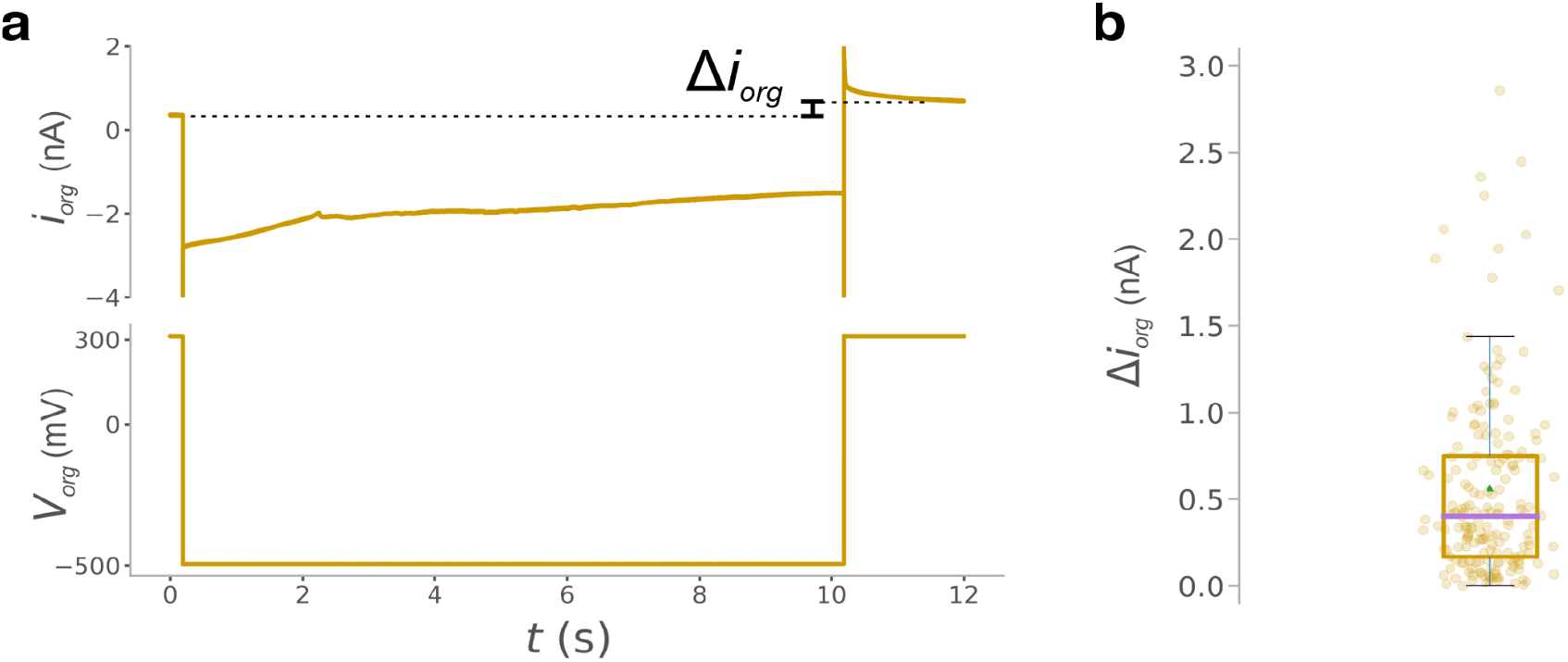
(A) Ion current *i*_org_ and applied voltage *V*_app_ trace in the organic barrel during nanobiopsy. After the extraction (t>10 s), the ion current *i*_org_ is characterised by a higher magnitude due to the cytoplasm entering the nanopipette which decreases the nanopipette resistance. (B) Boxplot showing the distribution of the increase in current magnitude Δ*i*_org_ (n=184).

This behaviour is driven by the velocity of the displacement of the liquid-liquid interface between the solution in the barrel (organic phase) and the cytoplasm (aqueous phase) which is fast at the beginning and decreases over time. At this point, the magnitude of the ion current is approximately equal to *i*_org_ ∼ 2.20 nA and it slowly decreases until reaching *i*_org_ = 0.70 nA, when the equilibrium is re-established. In the example trace, the increase in current magnitude due to ingress of cytoplasm after the extraction is equal to Δ*i*_org_ = 0.35 nA. **Figure 3b** shows the boxplot with the distribution of the current magnitude increase Δ*i*_org_ obtained after the extraction phase of 184 nanobiopsied cells with median value (purple line) equal to Δ*i*_org_ = 0.40 nA. The value Δ*i*_org_ can be used to estimate the cytoplasmic volume extracted^15^. Our estimation indicates 75% of samples showing an extracted volume ≤ 200 fl. However, this figure is only qualitative as the geometrical and electrical assumptions required for the estimation have large errors and cannot be used to precisely estimate the extracted volume. Nevertheless, **Fig. 3b** demonstrates the reproducibility of the protocol and indicates that it is very likely that a less than a pL is extracted from the cell. Further ion current traces recorded during nanobiopsy are shown in **Supplementary Fig. SF1.6**.

### SINGLE-CELL RNA SEQUENCING

Following nanobiopsy, the nanopipette is placed in a tube preloaded with 2 μl of lysis buffer containing an RNAse inhibitor to prevent mRNA degradation and immediately immersed in liquid nitrogen for short-term storage. The time elapsed from the nanopipette withdrawal to storage of the sample in liquid nitrogen was < 3 minutes and includes the time required to withdraw the nanopipette from the culture solution, break the nanopipette tip in the tube containing lysis buffer, label the tube and immerse it in liquid nitrogen. Next, the mRNA collected is reverse transcribed into cDNA and amplified according to Smart-Seq2^21^ and single-cell libraries are prepared for subsequent next-generation RNA sequencing and gene expression profiling.

The nanobiopsy technology allows the longitudinal gene expression profiling of single cells in culture. As a proof of concept, we investigated the transcriptional reprogramming in GBM cells in response to treatment. M059K GBM cells are amenable to nanobiopsy and can be visually tracked, along with their progeny, over a 72-hour time frame without leaving the field of view. Topographical mapping with SICM and fluorescent staining was performed on some of the M059K GBM cells to assess the average cell height, surface area and volume, this information was then used to guide the location of the nanobiopsy procedure (**Supplementary Fig. SF2.1**). For consistency, all nanobiopsies were carried out in the perinuclear region, and fluorescent staining indicated the presence of mitochondria and the endoplasmic reticulum in that region (**Supplementary Fig. SF2.2**). To enable longitudinal tracking of individual GBM cells over time, mixed cultures of GFP-transfected (M059K_GFP_) and wild type (M059K_WT_) cells were plated on gridded dishes at a ratio of 1:350 (M059K_GFP_ : M059K_WT_). This ratio facilitated the identification of individual M059K_GFP_ surrounded by M059K_WT_. The experimental setup and use of gridded dishes to locate cells, including following division, is outlined in **Supplementary Section S2.3**. In order to perform longitudinal analysis, it is crucial to ensure that the same cell is sampled. Therefore, the total (Δ*x*_tot_) and maximum (Δ*x*_max_) cellular migration over the planned 72-hour time course was quantified and showed not to differ between repeats (t-test, p>0.5) and to be substantially less (Δ*x*_max_ < 800 μm, Δ*x*_max_ < 600 μm) than the field of view (1300 × 1300 μm). Furthermore, the largest area enclosing all progeny, in cases where the cell divided over the time course, was 0.35 mm^2^ which is ∼5 times smaller than the area of the field of view (1.69 mm^2^) (**Supplementary Fig. SF2.3**).

We are therefore confident that the same cell was longitudinally sampled. Next, we tested whether the same M059K GBM cell could be sampled at different time points and we followed the same cell over the 72-hour time course. Results showed that the cell was successfully biopsied and injected on day 1, imaged on day 2 and day 3, and longitudinally biopsied and injected on day 4 (**Supplementary Fig. SF2.4**). Having established the suitability of the nanobiopsy for longitudinal sampling of the same GBM cell, we performed longitudinal sampling of M059K GBM cells through standard therapy as depicted in **Fig. 4a**. Single M059K_GFP_ cells were identified and their location recorded. A first nanobiopsy of the cell was executed to extract cytoplasm (day 1 sample) and inject a fluorescent dye. This was repeated on day 4 on the same cell, or each progeny thereof. On day 2, half of the dishes had been subjected to non-surgical elements of standard GBM treatment, including 2 Gy irradiation (IR) and 30 μM Temozolomide (TMZ), while the remaining half were left untreated. **Figure 4b** shows the optical micrographs of an individual M059K_GFP_ cell that was biopsied and injected on day 1, treated, and longitudinally biopsied on day 4 while **Fig. 4c** shows the case of an individual M059K_GFP_ that was biopsied and injected on day 1 and that divided following standard therapy and whose progeny was longitudinally biopsied and injected on day 4. Similar examples for the longitudinal samples collected from the untreated cells are shown in **Supplementary Fig. SF2.5**. The total number of nanobiopsies taken for each group (treated and untreated) at each timepoint (day 1 and day 4) is summarised in the schematics in **Fig. 4d**. 256 samples were collected from M059K_GFP_ cells of which 71 were longitudinal samples from 32 treated and 39 untreated cells. 12 treated and 12 untreated cells sampled longitudinally did not divide over the 72 hours while 7 untreated and 7 treated cells sampled longitudinally divided. When comparing between treated and untreated cells, there was no significant difference in the number of cells that underwent day 1 nanobiopsies and died versus survived (chi-squared, p=0.24), nor that survived and divided versus survived and didn’t divide (chi-squared, p=1.0). SmartSeq2 was performed to create sequencing libraries from the extracted cytoplasm samples, as well as from whole-cell lysates of treated and untreated M059K as a comparison (**Supplementary Section S2.6** and **Supplementary Fig. SF2.6**). Quality metrics of the two datasets (**Supplementary Fig. SF3.1**) were generally comparable, though the nanobiopsies had higher total reads (mean=6.5×10^6^ vs. 2.8×10^6^), fewer expressed genes (mean=497 vs. 1100), lower % of bases aligning to mRNA regions (mean=0.28 vs. 0.82), and higher % mitochondrial gene counts (mean=23.4 vs. 7.5). Both datasets were filtered for >150 expressed protein coding genes, <10% ribosomal bases and <30% mitochondrial gene counts, with the latter allowing for the high concentration of mitochondria in the nanobiopsied region. The sequencing metrics after filtering are shown in **Fig. 5a**. Gene expression profiles were generated for each nanobiopsy and whole cell sample and plotted using UMAP. Both the whole-cells and longitudinal nanobiopsies showed separation between treated and untreated samples (**Fig. 5b**), and this separation was not due to confounding from any observed technical biases (**Supplementary Fig. SF3.3**), suggesting that the nanobiopsies are capturing true biological effects of the treatment. Furthermore, by comparing day1 to day4 nanobiopsy positions on the UMAP plots we see differing trajectories of untreated and treated cells (**Fig. 5c**). It has been shown previously that GBM cells lie on an axis of proneural (PN) to mesenchymal (MES) cell subtypes, and that the distributions of these subtypes change through treatment^22^. It isn’t, however, known whether the change in distributions is due to differences in growth and death rates of each subtype or instead due to cells directly changing subtype through transcriptional reprogramming. To investigate this, we classified both treated and untreated M059K cells at each time point as either PN or MES (**Supplementary Fig. SF3.4**) by applying gene set variation analysis (GSVA) to the nanobiopsy gene expression profiles (**Fig. 5d**). There was no difference in the propensity of cell subtypes at day 1, irrespective of treatment, to subsequently die (Chi-squared, p=0.85) or survive (Fisher’s Exact Test, p=0.67). There was also no difference, irrespective of treatment, in the proportion of cell subtypes at day 4 (Fisher’s Exact Test, p=0.46). There was, however, a significant difference in the proportion of cells that switched subtype, or produced progeny with a different subtype when dividing, over time: of the cases where both day-1 and longitudinal biopsies had passed filtering, untreated cells switched subtype in 7/10 cases whereas treated cells only switched in 1/9 (**Supplementary Fig. SF3.4**, Fisher’s Exact Test, p=0.0094). This suggests that untreated cells are significantly more plastic over the 3-day time course than treated cells, which is further indicated in the changes in both PN (absolute change in score is 3.0-fold higher in untreated versus treated cells; T-test, p=0.041) and MES scores (absolute change in score is 2.6-fold higher in untreated versus treated cells; T-test, p=0.028) over time.

**Figure 4.**
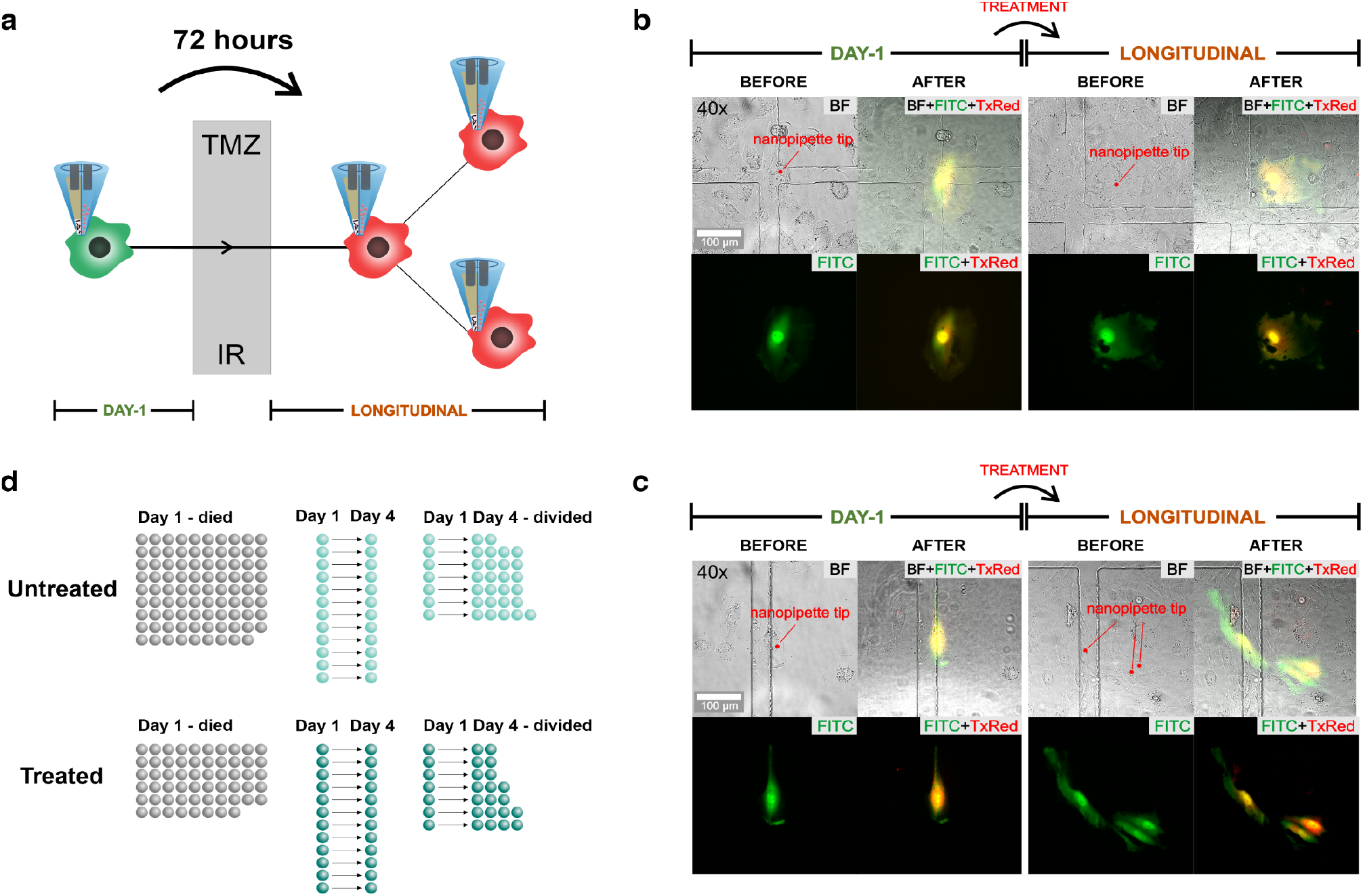
Longitudinal nanobiopsy of the same glioblastoma cells through standard therapy. (A) An individual GBM cell is nanobiopsied and injected (day-1) and standard treatment of temozolomide (TMZ) and irradiation (IR) is applied after which the same GBM cell or its progeny is nanobiopsied and injected a second time (longitudinal) 72 hours following the first nanobiopsy. (B) Optical (BF) and fluorescence (FITC, TxRed) micrographs of an individual M059K_GFP_ cell before and after nanobiopsy and nanoinjection and before (day-1) and following standard treatment (longitudinal). (C) Optical and fluorescence micrographs of an individual M059K_GFP_ cell that is nanobiopsied and nanoinjected on day 1, survive treatment and divide, whose progeny is nanobiospied and nanoinjected a second time following treatment. (D) Illustration of the nanobiopsy count of untreated (blue) and treated (red) cells that were nanobiopsied on day 1 and died (Day 1 – died), nanobiopsied on day 1, survived, didn’t divide and nanobiopsied again on day 4 (Day 1 Day 4) and cells that are nanobiopsied on day 1, divided, and whose progeny was nanobiopsied again on day 4 (Day 1 Day 4 – divided).

**Figure 5:**
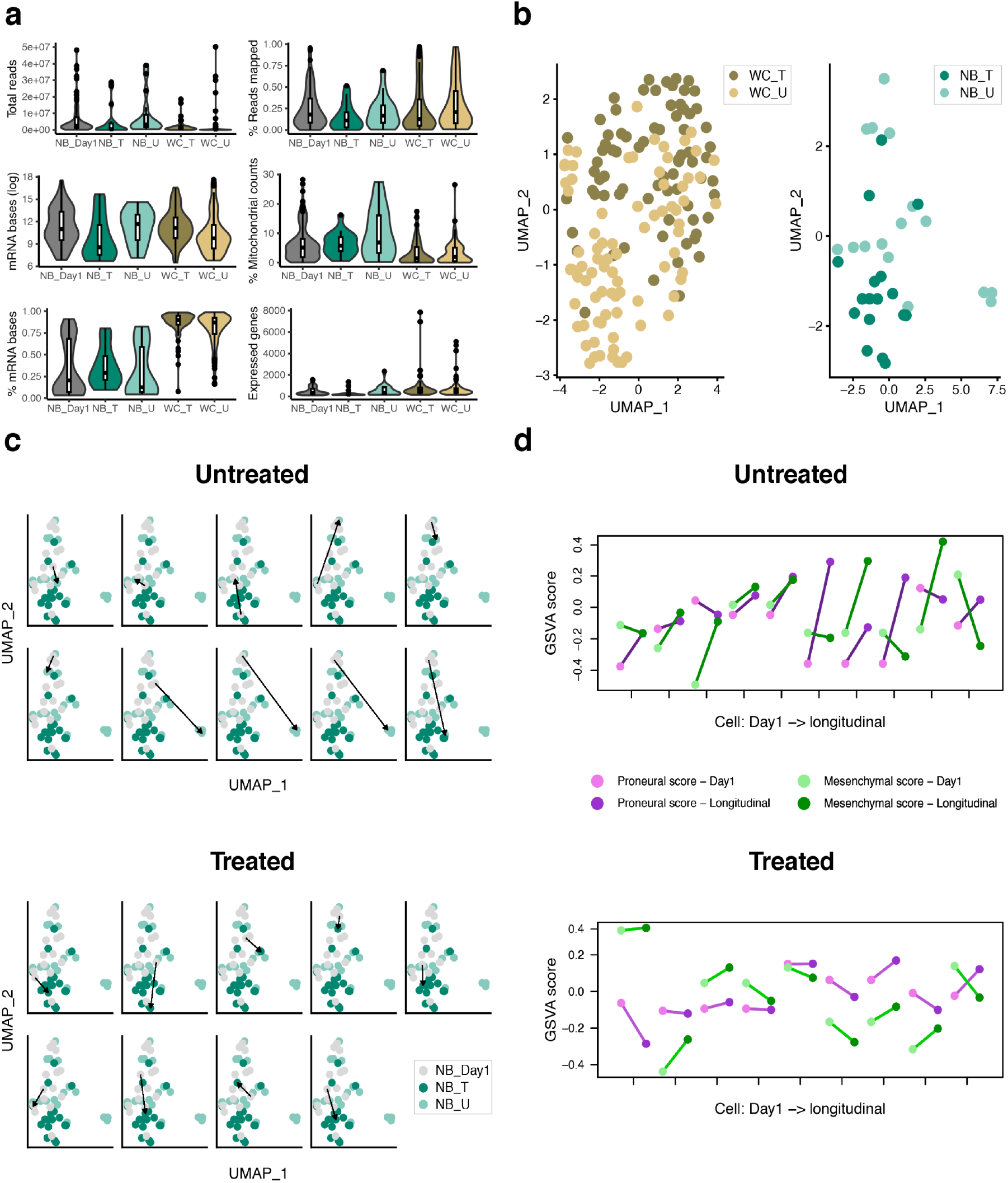
Data analysis of nanobiopsy and whole-cell lysate samples. a) Main sequencing metrics of day-1 nanobiopsy (NB_Day1) longitudinal nanobiopsy of treated (NB_T) and untreated (NB_U), and whole-cell lysate treated (WC_T) and untreated (WC_U) samples. b) UMAP visualisation of treated and untreated whole-cell lysate and longitudinal nanobiopsy samples. c) UMAP visualisation of paired Day-1, longitudinal treated and untreated nanobiopsy samples with arrows indicating the change in position of the paired day-1 and longitudinal sample. d) Cell scores from GSVA of proneural (purple) and mesenchymal (green) phenotypes for the paired day-1 (light colour) and longitudinal (dark colour) samples. The crossing of the lines indicates a subtype switch of the cell.

## DISCUSSION

There is significant interest in emerging technologies that can enhance our ability to understand and analyse transcriptome dynamics. Recent developments in *in vivo* fluorescence and super-resolution microscopy enabled the visualisation of transcription dynamics in living cells^23–25^. Single-cell RNA sequencing and lineage tracing have been coupled to combine clonal information with cell transcriptomes^26,27^, and metabolic labelling has been used to track newly synthesised RNA to allow the study of transcriptional dynamics to investigate how perturbations impact gene expression^28–30^. However, understanding how the initial state of a cell’s transcriptome influences its response to a perturbation remains challenging due to the assumptions required to infer a cell’s initial state when using methodologies which require cell lysis^31^. Recently, the development of Live-seq^6^ enabled single-cell profiling and functional analysis of the same cell at distinct time points, preserving cellular functions and viability. The development of such technologies is of crucial importance to complement already available high-throughput techniques to unravel the highly dynamic mechanisms of gene expression dynamics and phenotype variation which characterise healthy and diseased tissues.

In this study, we developed a SICM-based technology enabling the simultaneous nanoinjection of exogenous molecules into a living cell whilst extracting femtolitre-volumes of cytoplasm, via a double-barrel nanopipette. This method allows for the injection of a fluorescent dye into a cell to facilitate labelling and tracking and enables subsequent longitudinal nanobiopsy of the same cell to profile gene expression changes over time. Such investigations allow for unprecedented insights into how cells respond under different conditions or with different treatments and will likely be invaluable in many research settings including, for example, in understanding the development of treatment resistance in cancer. This may be especially relevant in GBM, where, following surgical resection and subsequent treatment with radiotherapy and chemotherapy, tumours almost always recur and are fatal, resulting in a median patient survival of just 15 months^32^. It is not known what allows the GBM cells to resist treatment, but recent studies suggest the involvement of transcriptional reprogramming and shifts to different cell states^33–36^. These investigations have largely relied on comparing cell populations from distinct pre- and post-treatment tumour samples, and therefore, only inferences can be made about the nature of any transcriptional reprogramming. We applied the nanobiopsy technique to M059K GBM cells to obtain sequential transcriptional profiles from the same cells as they undergo standard treatment, which includes radiotherapy and chemotherapy. The analysis of the sequencing metrics indicates that the nanobiopsy samples exhibit lower overall quality compared to the whole-cell lysate samples. This outcome was expected since the nanobiopsy samples represent only a small fraction of the total cytoplasmic mRNA, as supported by the estimated extracted volume (femtolitres) compared to the cytoplasmic volume (picolitres). Approximately 55% of the nanobiopsy samples met the filtering criteria. Of these, a separation between treated and untreated longitudinal nanobiopsy samples was observed on a UMAP plot. These results were similar to those obtained from whole-cell lysates, indicating that the mRNA extracted through nanobiopsy can indicate the cell’s transcriptional profile. The cell phenotype scores of paired day-1 and longitudinal samples revealed that treated cells tend to maintain the same phenotype during therapy, while untreated cells are more likely to switch transcriptional state over a 72-hour period, suggesting that treatment either induces or selects for higher transcriptional stability.

Future developments of the nanobiopsy technology will include increasing the extraction volume and optimising the protocol for the reverse transcription and library preparation to enhance the sequencing data quality, following what was done for the live-seq technique. We anticipate that the nanobiopsy technique will serve as a catalyst for the advancement of novel technologies in single-cell sampling, thereby expanding the rapidly evolving domain of longitudinal transcriptomics.

## METHODS

### CELL CULTURE

HeLa and M059K cells were cultured in Dulbecco’s Modified Eagle Medium/Nutrient Mixture F-12 (DMEM/F-12) (Gibco) supplemented with 10% Fetal Bovine Serum (Gibco) and 1x Penicillin/Streptomycin solution (Life Technologies) in a 5% CO_2_ humidified atmosphere at 37°C and maintained at less than 80% confluence before passaging. The GFP-transfected M059K cell line was generated using the Dharmacon GIPZ™ Lentiviral shRNA library encoding the green-fluorescent protein TurboGFP, denoted as M059K_GFP_.

### NANOPIPETTE FABRICATION

Double-barrel nanopipettes were pulled out of quartz theta capillaries (O.D. 1.2 mm, I.D. 0.9 mm, QT120-90-7.5, World Precision Instruments) using a CO_2_ laser puller (Sutter P2000, Sutter Instrument). A two-line program (Line 1: HEAT 750, FILAMENT 4, VELOCITY 30, DELAY 150, PULL 80 / Line 2: HEAT 680, FILAMENT 3, VELOCITY 40, DELAY 135, PULL 160) was used to generate double-barrel nanopipettes with pore dimension of the individual barrel ∼150 nm.

### SICM SETUP

The double-barrel nanopipette was mounted on a custom designed holder. An Ag/AgCl and an Ag electrode were inserted in the aqueous and organic barrel, respectively. Each electrode was connected to an headstage amplifier (Axopatch CV-7B) mounted on the SICM frame. The headstage amplifiers were connected to a patch-clamp amplifier (Multiclamp 700B) and to an analog-to-digital converter (Digidata 1550B). The ion current was sampled using a sampling frequency of 10 kHz. The SICM setup was comprised of an Axon MultiClamp 700B amplifier, MM-6 micropositioner (Sutter Instrument) and a P-753 Linear actuator (Physik Instrumente) to allow precise three-dimensional movement of the micropipette. The SICM software was used to control the positioning and topographical scanning capabilities of the SICM (ICAPPIC, London, UK). The z-piezo actuator had travel range 38 μm while the travel range for x and y-piezo was 96 μm. An Eclipse Ti2 inverted microscope (Nikon Instruments) and LED illumination system (pE-4000 CoolLED) with filter sets for DAPI, FITC, TxRed were used for bright field and epifluorescence imaging. An ORCA-Flash4.0 V3 Digital CMOS camera (C13440-20CU, Hamamatsu) was used to acquire optical and fluorescent micrographs.

### LONGITUDINAL NANOBIOPSIES

#### Oligos

<u>Oligo-dT30VN:</u> 5’ - AAGCAGTGGT ATCAACGCAG AGTACTTTTT TTTTTTTTTT TTTTTTTTTT TTTTTVN - 3’

<u>Template Switch Oligo (TSO):</u> 5’ - AAGCAGTGGTATCAACGCAG AGTACrGrG+G - 3’ where “r” indicates a ribonucleic acid base and “+” a locked nucleic acid base

<u>IS-PCR primers:</u> 5’ - AAGCAGTGGTATCAACGCAGAGT - 3’ All oligos were purchased from Integrated DNA Technologies.

#### Solutions

<u>Lysis buffer:</u> was prepared in aliquots of 2 μl stored at -80°C and contained 0.04 μl 10% Triton X-100 (Sigma), 0.1 μl SUPERasin RNAse Inhibitor (20 U/μl) (Thermo Fisher Scientific), 1.86 μl Nuclease Free Water, for a total volume of 2 μl.

<u>Mastermix 1 (Priming mix)</u> was prepared in aliquots of 19 μl (10 rxs) and stored at -80°C and contained 1 μl dNTP mix (10 mM each, Thermo Fisher Scientific), 0.9 μl Nuclease Free Water (Thermo Fisher Scientific) for a total volume of 1.9 μl per reaction. Prior to use, 1 μl of Oligo-dT30VN (100 μM) was added to the thawed 19 μl aliquot for a total volume of 20 μl (10 rxs).

<u>Mastermix 2 (RT-PCR mix)</u> was prepared in aliquots of 48.5 μl (10 rxs) and store at -80°C and contained 0.06 μl MgCl_2_ (1M, Sigma), 2 μl Betaine (5M, Sigma), 0.5 μl DTT (100 mM, Thermo Fisher Scientific), 2 μl 5x Superscript first strand buffer (Thermo Fisher Scientific), 0.29 μl Nuclease Free Water (Thermo Fisher Scientific) per reaction. Prior to use, 1 μl TSO oligo (100 μM), 2.5 μl SUPERasin RNAse Inhibitor (20 U/ μl) and 5 μl Superscript II (200 U/ μl) were added to the thawed 48.5 μl aliquot for a total of 57 μl (10 rxs).

<u>Mastermix 3 (IS-PCR)</u> was freshly prepared and contained 12.5 μl Kapa HiFi Hotstart Readymix (2x, Roche), 0.25 μl IS PCR Primers (10 μM) and 2.25 μl Nuclease Free Water for a total of 15 μl per reaction.

#### Cell preparation

M059K_GFP_ and M059K_WT_ cells were detached from the flasks using Trysin-EDTA solution (Sigma Aldrich) and centrifuged at 1000 RCF for 5 minutes. Cells were resuspended in 1 ml of fresh medium and counted using a Hemocytometer (DHC-B01-50, Burker). M059K_GFP_ and M059K_WT_ cells were diluted to the same concentration and seeded into a 35-mm μ-Dish, high Grid-500 (IBIDI IB-81166) with seeding density 2.5 · 10^4^ cells/dish at a ratio of 1:350 (M059K_GFP_ : M059K_WT_) to facilitate the identification and tracking of an individual M059K_GFP_ cell on the grid. After seeding, cells were incubated in a 5% CO_2_ humidified atmosphere at 37°C for 12-18 hours before nanobiospy. Shortly before the experiment, the culture medium was replaced with 2 ml of phenol free DMEM/F-12 (Gibco) supplemented with 10% Fetal Bovine Serum (Gibco) and 1x Penicillin/Streptomycin solution (Life Technologies).

#### Probe preparation

One barrel of the nanopipette was filled with 5 μl of 100 μM fluorophore (ATTO 488, 41051 Sigma Aldrich, for HeLa and ATTO 565, 75784 Sigma Aldrich, for M059K) in 0.1 M KCl (Sigma Aldrich). The operation was performed ensuring that dry conditions were maintained to avoid water to wet the backend of the nanopipette which represents the main cause of crosstalk between the two barrels. The second barrel was filled with 5 μl of the organic phase mixture of 10 mM THATPBCl in 1,2 DCE (Sigma), following silanisation of the backend of the nanopipette by exposure to Trichloro(1H, 1H, 2H, 2H-perfluorooctyl)silane (Sigma Aldrich) for 10 seconds until the glass wall appeared mat. This operation makes the glass hydrophobic and reduce the insurgence of crosstalk during the experiment. After filling the two barrels, the nanopipette was left for 5 minutes at 60°C to ensure that the back of the nanopipette was dry before the experiment. The ion current generated in the aqueous barrel was used as SICM feedback to drive the nanopipette towards and inside the cell. The electrolyte solution in the aqueous barrel has high electrical conductivity (σ_aq_ = 1.3 S/m) and generates an ion current with greater magnitude than the one generated in the organic barrel, where the conductivity of the organic solution is two orders of magnitude smaller (σ_org_= 0.011 S/m)^15^.

#### Nanobiopsy

An individual M059K_GFP_ was identified on the grid using a 10x magnification objective (Nikon) in epifluorescence microscopy, ensuring that no other M059K_GFP_ were detected in the same field of view. Next, a 40x objective (Nikon) was used to acquire the fluorescence and optical micrograph of the cell prior nanobiopsy. The double-barrel nanopipette was mounted on the SICM holder and was immersed using the linear actuator of the SICM until an ion current through the aqueous barrel was detected. During immersion, the potential applied to the electrode in the aqueous and organic barrel was *V*_aq_ = 200 mV and *V*_org_ = 300 mV, respectively, with respect to a reference Ag/AgCl electrode immersed in the culture media. In the approach phase, the SICM working in hopping mode (HPICM)^17^ moved the double-barrel nanopipette towards the cell membrane until the setpoint current (99.5% of reference current) was detected in the aqueous barrel. At this point the system stopped the nanopipette and retracted it to a distance from the cell equal to the hopping height parameter that was set to 2 μm. Following approach, the double-barrel nanopipette was visualised at the microscope and it was manually positioned onto the perinuclear region of the cell by means of a micromanipulator. In the nanoinjection phase, the double-barrel nanopipette was first moved vertically towards the cell of a distance equal to the hopping height (2 μm) to reach a position which is estimated to be <500 nm from the cell membrane. At this point, the ion current in the aqueous barrel dropped of 2% from the reference value because of the restricted ion flow due to the proximity of the membrane. Next, the potential applied to the electrode in the aqueous barrel was switched to *V*_aq_ = -500 mV to release the anionic fluorophore and the double-barrel nanopipette was moved towards the cell vertically with 100 nm steps at 100 nm/ms until the detection of the vibrational noise due to the proximity of the double-barrel nanopipette to the polymer coverslip of the dish. Upon detection of the vibrational noise, the indentation of the nanopipette to the cell membrane is enough to puncture the membrane and access the cytoplasm. Next, the nanopipette was withdrawn of 100 nm and was kept in position for ∼60 s to allow the electrophoretic release of the anionic fluorophore into the cytoplasm. At the end of the nanoinjection phase, the cell emitted a red-fluorescent signal that was visualised in epifluorescence. Following nanoinjection, the cytoplasmic extraction is enabled by switching the potential applied to the electrode in the organic barrel to *V*_org_ = -500 mV for 10 s. The application of the negative potential generated an inflow of cytoplasm in the nanopipette via electrowetting of the organic phase in the nanopipette barrel. A successful cytoplasmic extraction resulted in an increased magnitude of the ion current in the organic barrel *i*_org_ due to the ingress of cytoplasm whose conductivity (σ_cytoplasm_∼0.35-0.5 S/m)^37^ is 30-45 times greater than the conductivity of the organic solution (σ_org_= 0.011 S/m). Following cytoplasmic extraction, the double-barrel nanopipette was withdrawn and unloaded from the SICM and immediately placed in a tube preloaded with 2 μl of lysis buffer containing an RNAse inhibitor to prevent mRNA degradation. The tube was immersed in liquid nitrogen and stored until reverse transcription and cDNA amplification via Smart-Seq2 which was performed in all cases <7 days following nanobiopsy. The time elapsed from the withdrawal of the nanopipette to the immersion of the tube in liquid nitrogen was estimated to be < 3 minutes, including the time required to withdraw the nanopipette from the culture solution (∼ 30 s), remove the electrode and unload the nanopipette (∼ 30 s), gently break the tip in the tube containing lysis buffer (∼30 s). label the tube (1 min) and immerse the tube in liquid nitrogen (<30 s). Following nanobiopsy, the dish containing the biopsied cells was left to recover in a 5% CO_2_ humidified atmosphere at 37°C for 24 hours. On day 2, the dish was either subjected to non-surgical elements of standard GBM treatment consisting of 2 Gy irradiation and 30 μM Temozolomide (treated samples) or to mock irradiation (untreated samples). On day 3, cells were allowed to recover in the incubator. On day 4, the same cell or the progeny thereof was localised in the same location on the grid in epifluorescence and bright field microscopy and the same nanobiopsy procedure was carried out to perform longitudinal cytoplasmic extractions.

#### Reverse transcription and cDNA amplification

The Smart-seq2 protocol^21^ was adjusted to optimise the reverse transcription and cDNA amplification of the mRNA extracted via nanobiopsy. An aliquot of Mastermix 1 and Mastermix 2 were thawed on ice and the required reagents were added to complete the reaction mix. A volume of 2 μl of Mastermix 1 was dispensed in the PCR tube containing the nanobiopsy sample in 2 μl lysis buffer. Next, the tube was vortexed and briefly centrifuged. The tube was incubated on a thermocycler (Eppendorf Mastercycler Nexus GX2) at 72°C for 3 minutes to anneal the Oligo-dT30VN to the poly-A tail of the mRNA transcripts. After incubation, the tube was immediately placed on ice for 1 minute. Next, the tube was centrifuged for few seconds to collect condensations. A volume of 5.5 μl of Mastermix 2 was dispensed into the PCR tube. Next, the tube was vortexed and briefly centrifuged. The total reaction volume was ∼10 μl. The following thermal cycling program was performed using a thermocycler: (1) 42°C for 90 min; (2) 50°C for 2 min; (3) 42°C for 2 min, go to step (2) for 10 cycles; (4) 70°C for 15 min and (5) 4°C hold. The cDNA resulted from the reaction was amplified using a hot-start PCR to obtain the concentration required for library preparation. 15 μl of Mastermix 3 were added to the tube containing the RT-PCR product for a total reaction volume of ∼25 μl. The following thermal cycling program was performed using a thermocycler. (1) 98°C for 3 min; (2) 98°C for 20 s; (3) 67°C for 15 s; (4) 72°C for 6 min, go to step (2) for 26 cycles; (5) 72°C for 5 min and (6) 4°C hold. The amplified cDNA was purified using the HighPrep PCR Clean-up System (Sigma Aldrich AC-60050) adding a volume of magnetic beads equal to 0.8x the sample volume. The individual cDNA samples were quantified using the fluorometric assay QuantiFluor dsDNA System (Promega, E2670) using a 500 bp dsDNA calibration sample. The quality and fragment size of the amplified cDNA was quality checked using a D5000 tape (Agilent).

#### Library preparation and Next-Generation Sequencing

Next-generation sequencing libraries were prepared using the Nextera XT DNA Library Preparation Kit (FC-131-1096, Illumina). Following fluorometric quantification, each cDNA sample was diluted to a concentration of 0.2 ng/μl, which is the concentration needed for the tagmentation reaction. cDNA samples were diluted in a 96-well plate (Plate 1: cDNA). The concentration of 12 wells (diagonal) of the plate containing the diluted cDNA was checked using the QuantiFluor dsDNA System (Promega, E2670) fluorometric assay as the initial cDNA concentration is a critical step to obtain libraries with adequate fragment length. The tagmentation mastermix consisting of 300 μl Tagment DNA Buffer and 150 μl Amplicon Tagment Mix for a total volume of 450 μl was prepared on ice and a volume of 3.75 μl of the tagmentation mastermix was dispensed into each well of a new 96-well plate (plate 2: tagmentation) on ice. Next, 1.25 μl of the diluted 0.2 ng/μl cDNA was transferred from Plate 1 to Plate 2. Plate 2 was sealed and centrifuged at 1000 g for 1 minute at 4°C and the tagmentation reaction was performed using a thermocycler: (1) 55°C for 10 min and (2) 10°C hold. Next, 1.25 μl of Buffer NT and 3.75 μl of Nextera PCR Mastermix (NPM) from the Illumina kit were added to each well of the 96-well plate on ice. For each well, a total of 2.5 μl of Illumina indexes (i5 index + i7 index) were added at a 1:1 ratio. Following indexing, the amplification of the tagmented DNA fragments was performed using a thermocycler: (1) 72°C for 3 min; (2) 95°C for 30 s; (3) 95°C for 15 s; (4) 55°C for 10 s, go to step (3) for 12 cycles; (5) 72°C for 5 min and (6) 4°C hold. At the end of the PCR reaction, the cDNA fragments are of ideal size and uniquely barcoded (library). The concentration of the individual libraries was spot checked on the diagonal of the 96-well plate (12 samples) to ensure that the library concentrations were not too different from each other. A multiplexed library was generated by transferring 5 μl of each individual library into a tube and the multiplexed library was purified using the HighPrep PCR Clean-up System using a library:bead ratio of 1:0.6. The size of fragments of the multiplexed library was checked on a D1000 tape (Agilent). All the multiplexed libraries generated were sequenced using NextSeq2000 (Illumina), a high-throughput flow cell and adopting a P2 200 cycle (100 Pair End) and P3 200 cycle (100 Pair End) strategy.

## WHOLE CELL LYSATE PREPARATION

The M059K_GFP_ cells were grown to 80% confluency inside tissue culture flask, the cells were then separated into two flasks by depositing the same number of cells and allowed to grow 50% confluency in both flasks before the next step. One of the two flask was then subjected to a treatment of 2 Gy irradiation and 30 μM Temozolomide, these cells were labelled as treated cells, the other flask of cells were labelled as the untreated cells. Both flasks were left inside the incubator for 24 hours prior to Fluorescence-Activated Cell Sorting (FACS). FACS were used to single cell sort the viable M059K_GFP_ into 96 wells plate. 2 μl of lysis buffer consists of 0.2% Triton X-100 and 1 U/μl SUPERasin RNAse inhibitor (Thermo Fisher Scientific) was added to all the wells of the 96 well plate, this procedure was carried out inside a RNAse decontaminated laminar flow hood. Prior to FACS, the treated cells were harvested and washed twice with ice cold PBS at the concentration of 0.5×10^6^ cells/ml and stained with DAPI (Sigma-Aldrich) at the concentration of 0.1 μg/ml to exclude dead cells. For FACS, the gates were adjusted to exclude cell debris and doublets by size and scatter selection, only the viable GFP positive cells were selected, and subsequently single cell sorted into the lysis buffer containing 96 wells plate. The sorted plates were immediately sealed, and centrifuged at low speed for 1 minute, snap froze with dry ice and stored at -80°C. The same harvesting and FACS procedure were used again to single cell sort the untreated cells on the same day. The plates were used within 2 weeks after the sorting. The single cell sorted plates were subjected to the adjusted Smart-Seq2 protocol^21^ as outlined in the longitudinal nanobiopsy procedure with the exception that all the reaction solutions were freshly prepared prior to procedure. The amplified cDNA samples were then used for library preparation as outlined in the longitudinal nanobiopsy procedure above.

### DATA ANALYSIS

Reads were trimmed with cutadapt (v4.1) to remove Nextera adapters and low-quality bases with the parameters “-a CTGTCTCTTATA -A CTGTCTCTTATA --minimum-length 30 --overlap 5 -q 10”. Reads were then aligned and transcripts quantified using the GRCh38 genome with Gencode v27 basic annotations via STARsolo (v2.7.10b) with “-outFilterMatchNmin” set to 60 to ensure high stringency alignments.

Resulting BAM files were split using samtools split (v1.16.1) and sequencing metrics generated via Picard CollectRnaSeqMetrics (v2.20.2-SNAPSHOT). Nanobiopsies and whole cell lysates datasets were filtered for >150 expressed genes, <30% mRNA bases, and <10% ribosomal bases. Genes were filtered separately in each dataset for those that have >3 counts in each of 2 or more cells. Non-protein coding genes were also removed. Gene counts were normalised and scaled via Seurat (v4.3.0) using default settings with the addition of “scale.max = 5” and regressing out number of genes expressed and % mRNA bases, for the nano biopsies, and number of genes expressed for the whole cell lysates. The top 600 highly variable genes were identified in each using the Seurat “disp” method and used to perform PCA. UMAPs were then generated from the first 8 principal components. GSVA (v1.42.0) was run on the nanobiopsy scaled gene expression. Gene sets for mesenchymal and proneural phenotypes were taken from the top and bottom 50 weighted genes from PC1 for the 10x single-cell dataset in Wang et al.^22^. The stemness gene set was taken from Patel et al.^38^.

## DATA AVAILABILITY

The data that support the findings of this study are openly available from the University of Leeds data repository.

## CODE AVAILABILITY

The raw sequencing data and the code used for data analysis are openly available from the University of Leeds data repository and from https://github.com/GliomaGenomics/GBM_NB.

## ACKNOWLEDGEMENTS

This work was made possible owing to funding from The Brain Tumour Charity (TBTC) grant number GN-000482 (to LFS and PA); UK Research and Innovation (UKRI) grant number MR/T020504/1 (to LFS); European Commission H2020 grant number 812398 (to PA). This work was undertaken on ARC3, part of the High-Performance Computing facilities at the University of Leeds, UK. We thank Dr. Alexander Kulak for the help provided with SEM imaging of the double-barrel nanopipette.

## AUTHOR CONTRIBUTIONS

F.M. designed and performed all experiments and contributed to data analysis. C.C. designed and performed experiments and G.T. led and performed data analysis. L.F.S. and P.A. conceived the project, helped design the experiments and supervised the research. M.E., M.F., S.A., helped design the experiments and data analysis. M.R., C.L., I.C. provided technical help with library preparation and performed next-generation sequencing. I.M. provided training on Smart-Seq2 and helped with data analysis. All authors wrote and corrected the manuscript.

