## Supporting Information for "Single Cell Track and Trace: live cell labelling and temporal transcriptomics via nanobiopsy"

#These authors contributed equally

### **S1: Single-cell nanobiopsy technique**

#### *S1.1 Double barrel nanopipette fabrication and scanning electron microscopy (SEM).*

Double-barrel nanopipettes used in this work were fabricated frp, theta-quartz capillaries using a Sutter P-2000 laser puller using the following parameters:

|  | **HEAT** | **FILAMENT** | **VELOCITY** | **DELAY** | **PULL** |
| --- | --- | --- | --- | --- | --- |
| **LINE 1** | 750 | 4 | 30 | 150 | 80 |
| **LINE 2** | 680 | 3 | 40 | 135 | 160 |

The SEM images and the measured dimension of the individual barrels for four different nanopipettes in **FIGURE SF1.1** show that one barrel (barrel 1) is characterised by a bigger aperture than the aperture of the other barrel (barrel 2) which is due to the asymmetric pulling process of quartz-theta capillaries.

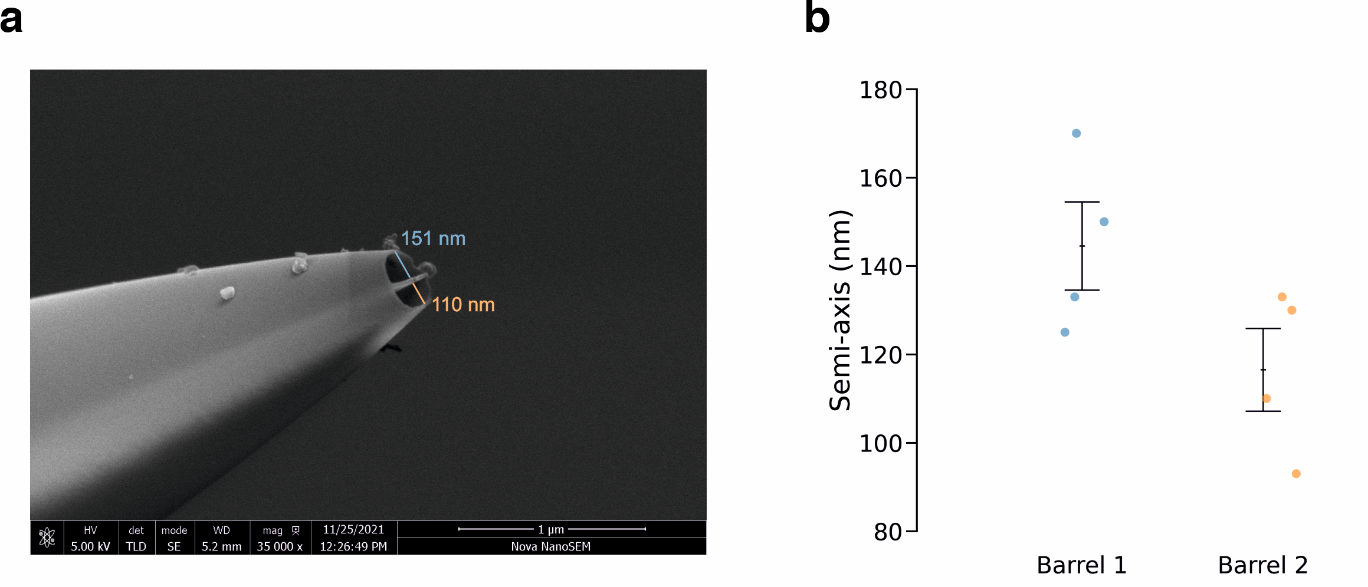

**Figure SF1.1.** (a) Scanning electron microscopy (SEM) micrograph showing the double-barrel nanopipette used in this work and (b) scatter plot showing the dimension of the semi-axis of each barrel across four different nanopipettes where barrel 1 is the big barrel and barrel 2 is the smaller one. The centre point indicates the mean, and the whiskers represent the standard error of the mean.

#### *S1.2 Electrical resistance of the aqueous and organic barrel*

The electrical resistance of the barrel filled with the aqueous solution of 0.1 M KCl and 100 µM ATTO 565 and the one filled with the organic solution of 10 mM THATPBCl in 1,2 DCE was calculated from the current trace of the aqueous and organic barrel using Ohm’s law. For the aqueous barrel (**FIGURE SF1.2a**), the median value for the electrical resistance was 45.7 MΩ. For the organic barrel (**FIGURE SF1.2b**), the median value for the resistance was 2.7 GΩ which is ~59 times higher than the resistance of the aqueous barrel. In both cases, the total number of data points is 256.

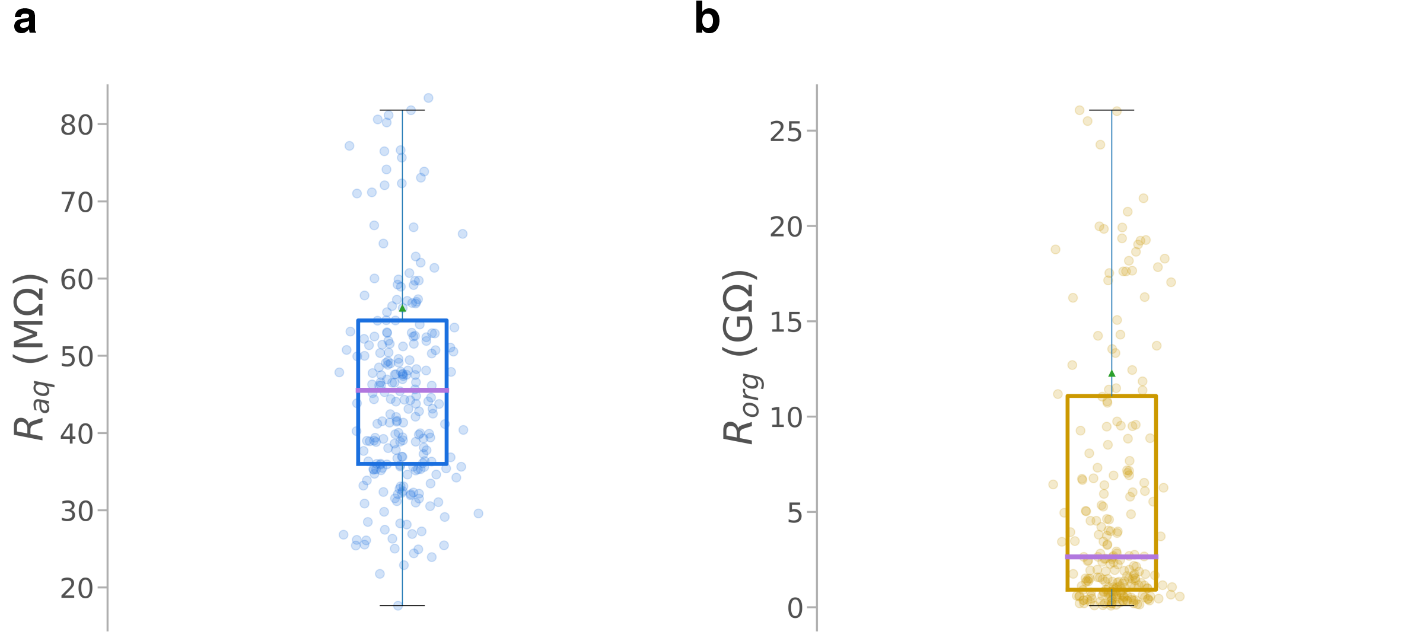

**Figure SF1.2.** Experimental electrical resistance of the aqueous (a) and organic (b) barrel of the nanopipettes used in the nanobiopsy experiments. Note the unit for *R*_org_ is GΩ. The green triangle and the purple line show the mean and median value, respectively. N = 256.

#### *S1.3 Example of electric signals recorded during the approach phase.*

In the approach phase (**FIGURE SF1.3a**), the double-barrel nanopipette automatically approaches the cell membrane driven by an SICM working in hopping mode with value of the setpoint current set to 99.5% of the baseline current and hopping height set to 2 µm. The approach of the cellular membrane is concluded when the current signature of the hopping mode shown in **FIGURE SF1.3b** is detected in the aqueous barrel, where the current *i*_aq_ drops to the setpoint current when the piezo actuator *z* reaches the hopping height value. At the end of the approach phase, the nanopipette is visible as a tiny bright dot in the field of view and it is manually moved to the biopsy location on the cellular membrane using a X-Y micromanipulator (**FIGURE SF1.3c-d**).

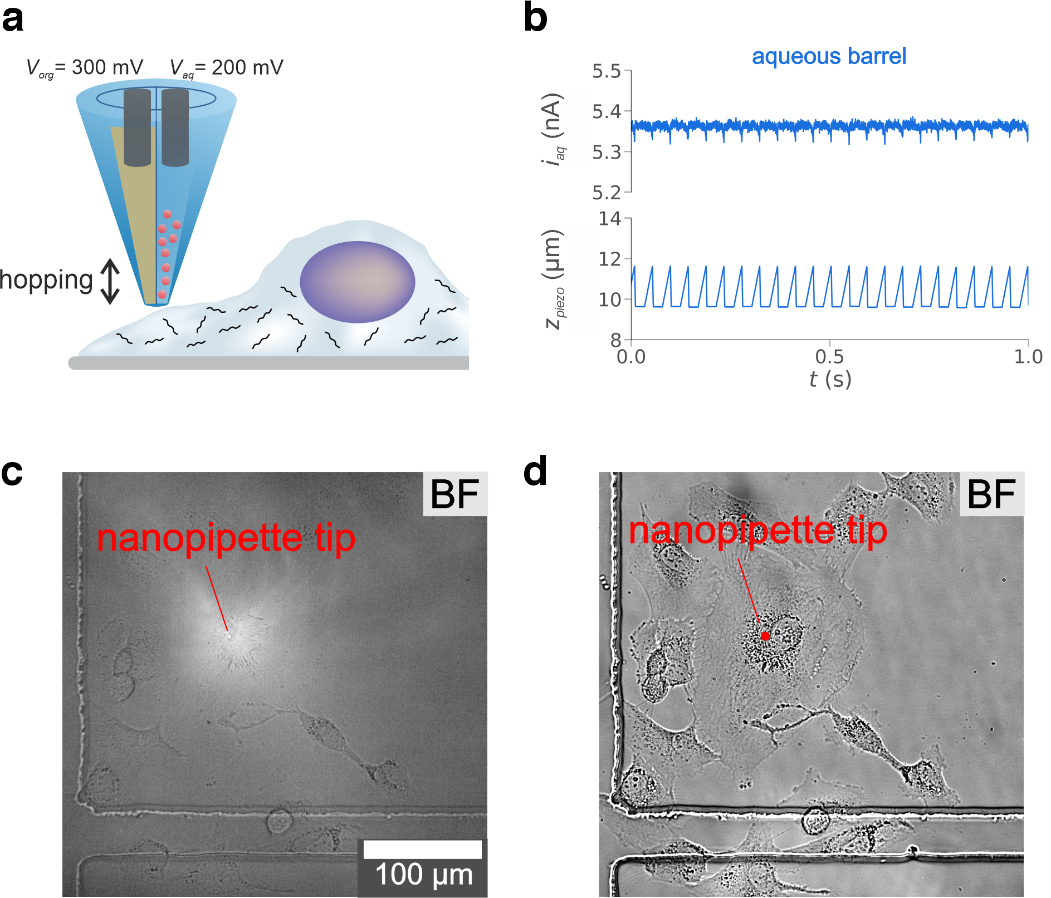

**Figure SF1.3.** Illustration of a double-barrel nanopipette approaching the cellular membrane (a) and experimental recordings of the ion current in the aqueous barrel i_aq_ and piezo vertical position z_piezo_ at the end of the approach phase (b), when the nanopipette tip appears as a bright dot under bright field (BF) microscopy (c). The position of the nanopipette after nanopipette withdrawal is marked with a red dot (d).

#### *S1.4 Power spectral density of the ion current in the nanoinjection phase.*

The signal for the ion current recorded in the aqueous barrel shown in **FIGURE a-b (main text)** can be analysed in the frequency domain to provide information on all the three sub-phases of the cell membrane and nanoinjection phase of the nanobiopsy experiment. **FIGURE S1.4** shows the spectrogram of the ion current trace *i*_aq_ shown in **FIGURE 2a-b (main text)** with the z-piezo trace plotted at the top (the z-piezo signal was flipped for ease of representation). The hopping phase (1) can be identified by the high power of the frequencies multiple of 25 Hz (hopping frequency) in the first second of the map. Next, at t∼5 s, a sudden increase in the power of all frequencies is observable upon the motion of the nanopipette from the hopping height (2 µm) to the surface of the cell (membrane touch phase (2)). The noise increase in the lower frequencies (<1000) is due to the contact of the nanopipette with the plasma membrane. Next, the potential to the electrode in the aqueous barrel is switched to -500 mV and a spike in all frequencies can be observed around t=10 s. After the potential switch, the nanopipette is lowered down with 100-nm steps until the vibrational noise (3) is observable as an increase in the power across all frequencies around t=38 s. At the point where the piezo reaches the maximum extension (t=46 s), the power of all frequencies decreases of several orders of magnitude for approximately 3 seconds. The decrease in the power is due to the ion current having magnitude equal to 0 in this time interval, due to the contact between the nanopipette and the surface of the Petri dish. This is the point of maximum indentation. Upon the nanopipette retraction of 100 nm, the ion current is re-established, and the spectrogram shows an increase in the power of all frequencies. Interestingly, after nanopipette retraction, the current continues to be characterized by higher power across all the spectrum (higher noise) that might be due to the cell cytoplasm which represents a noisier environment than the cell medium. The spectrogram of the ion current in the aqueous barrel of the nanopipette provides a summary of the cell membrane and nanoinjection phase. Compared to the time-domain signal, the spectrogram summarises all the information in a map that can be used in a post-processing to extract and compare the parameters describing the experiment. A post-processing analysis of these maps could help to identify the marker parameters that could distinguish a successful from an unsuccessful nanobiopsy. For example, the spectrogram in **FIGURE SF1.5a** shows the three independent phases for the membrane penetration and nanoinjection resulting in the injected cell emitting a red-fluorescent signal as shown in **FIGURE SF1.5b.** As opposite, the spectrogram in **FIGURE S1.5c** shows an irregular pattern with some intervals with no power in all frequencies because of the current being out of range of the analog-to-digital converter, resulting in the cell emitting no red fluorescent signal at the end of the procedure, as shown in **FIGURE S1.5d,** suggesting that the nanopipette failed to penetrate the cellular membrane.

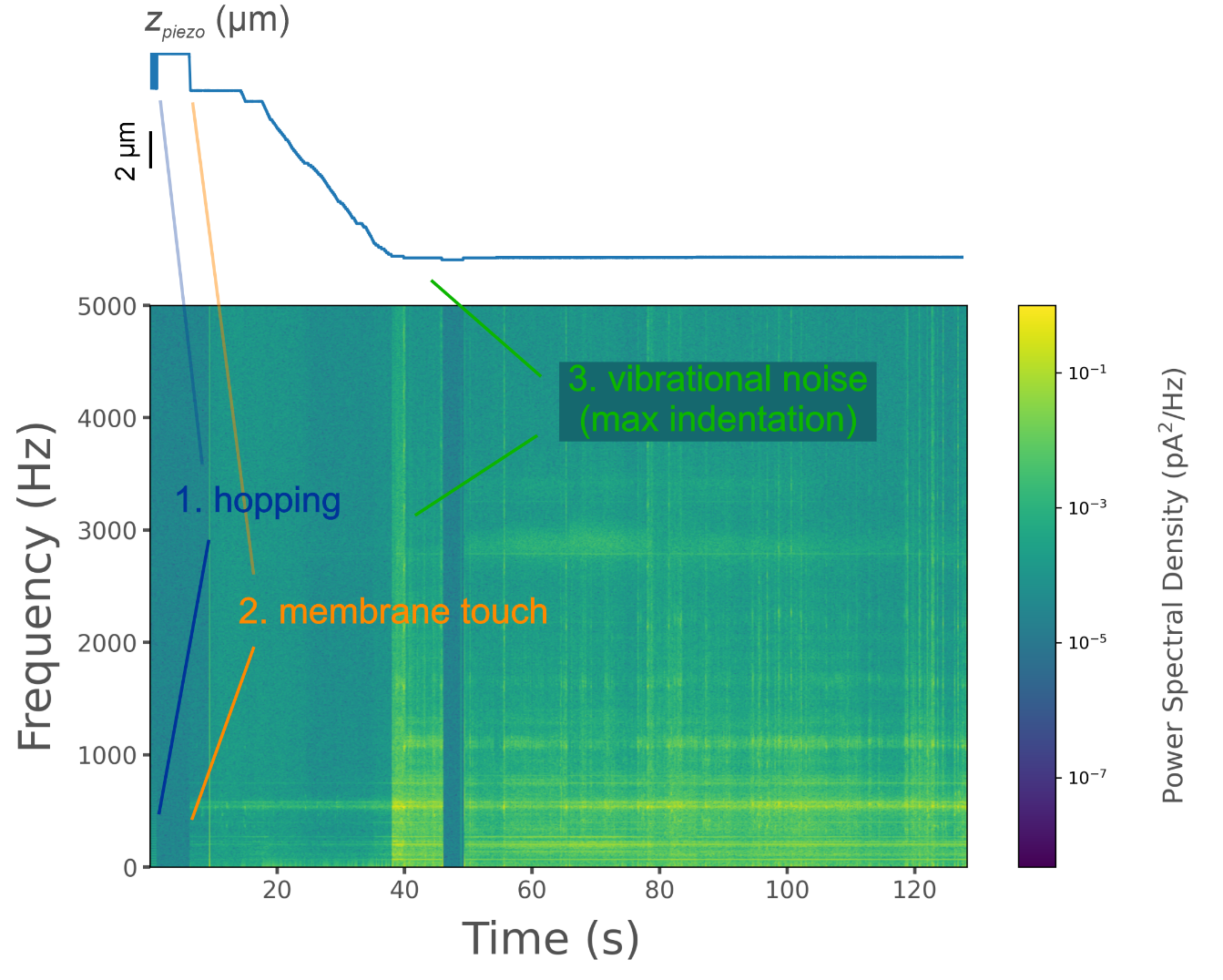

**Figure SF1.4.** Spectrogram for the ion current recorded during the membrane penetration and nanoinjection phase in a M059K cell. The z-piezo trace is shown at the top for reference and it was flipped to get a decrease in the signal when the nanopipette is lowered towards the cell cytoplasm. The power spectral density (pA2/Hz) is shown by the colour map for each frequency over time. A frequency signature identifies the three main phases of the cell penetration and injection experiment including hopping (1), membrane touch (2) and vibrational noise (3). The spectrogram was generated calculating the Fast Fourier Transform using Hamming window with length equal to 2048 samples (corresponding to 0.2 s) and an overlap of 256 samples. For a signal sampled with 10 kHz sampling frequency, the parameter used generate a spectrogram with time resolution 0.2 s and frequency resolution 5 Hz.

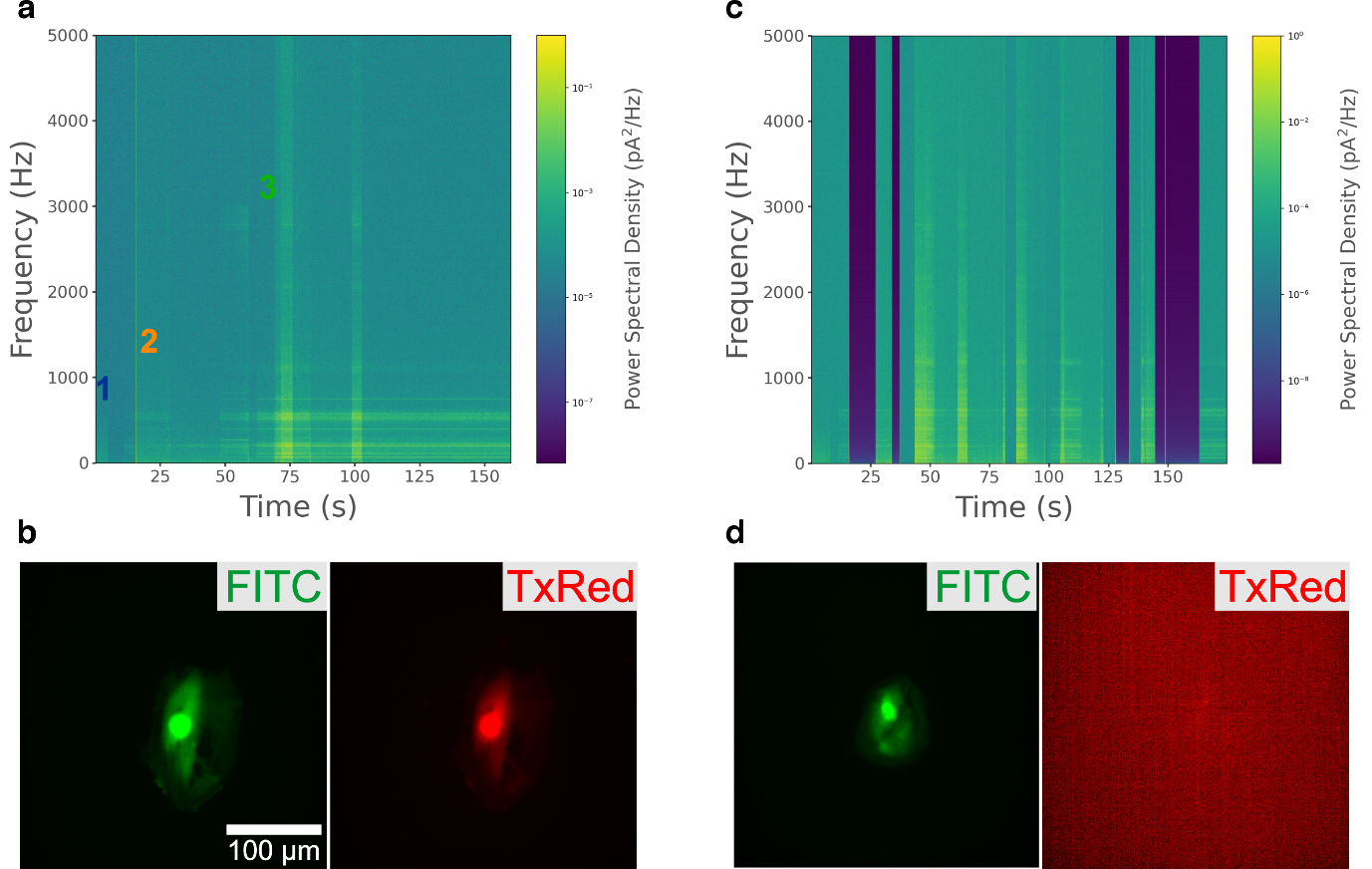

**Figure SF1.5.** Example of a spectrogram of the ion current signal in case of a successful (a) and unsuccessful (c) cell membrane penetration and nanoinjection. A spectrogram where all three sub-phases (i.e. hopping 1, membrane touch 2, vibrational noise 3) can be identified (a) results in the cell emitting a red-fluorescent signal following nanoinjection of a red fluorophore (b). A spectrogram with irregular pattern where the individual sub-phases cannot be identified (c) results in the cell emitting no red fluorescent signal at the end of the procedure (d).

#### *S1.5 Reproducibility of experimental traces during the nanoinjection phase*

The spectrogram for the ion current in the aqueous barrel *i*_aq_ during the nanoinjection phase can be used as quality control to check whether the nanopipette was successful at penetrating the cellular membrane and accessing the cytoplasm. **FIGURE SF1.6** shows some representative spectrograms recorded during the nanoinjection phase in the nanobiopsy of M059K_GFP_ GBM cells where the three subphases, namely hopping (1), membrane touch (2) and vibrational noise (3), were detected and the red fluorescent molecule was successfully injected as observable in the optical micrographs.

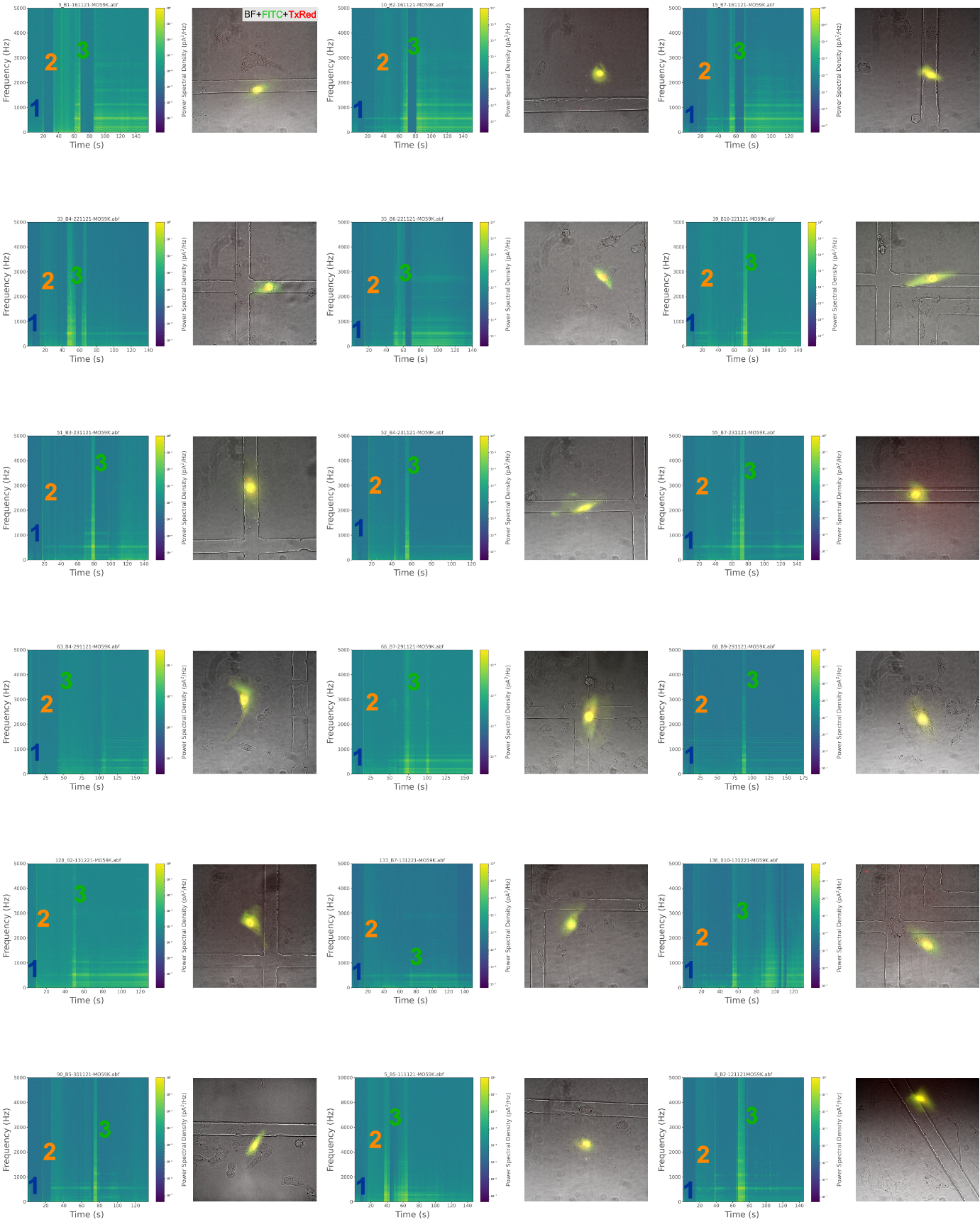

**Figure SF1.5.** Representative spectrograms of the ion current in the aqueous barrel i_aq_ recorded during the nanoinjection phase and optical micrograph of the cell after nanoinjection under false colour obtained by merging the bright field (BF), green (FITC) and red (TxRed) images. The three individual sub-phases hopping (1), membrane touch (2) and vibrational noise (3) can be identified in all spectrograms and can be used as an indication that the cellular cytoplasm was successfully accessed, as confirmed by the red fluorescent signal emitted by the cell after the procedure. Spectrograms were calculated from the ion current in the aqueous barrel i_aq_ recorded during 18 M059K_GFP_ nanobiopsies performed on 8 different days. Each nanobiopsy was performed using a different nanopipette.

#### *S1.5 Reproducibility of experimental traces during the nanobiopsy phase*

The trace for the ion current in the organic barrel *i*_org_ can be used to assess whether a cytoplasmic volume was extracted in the nanobiopsy phase. If the value for *i*_org_ after the potential switch *V*_org_=-500 mV for 10 s is greater than the value before the potential switch, a cytoplasmic volume is extracted. The ingress of cytoplasm in the nanopipette decreases the nanopipette resistance thus increasing the ion current *i*_org_ in the organic phase. **FIGURE SF1.6** shows the ion current *i*_org_ recorded from 24 randomly selected nanobiopsies.

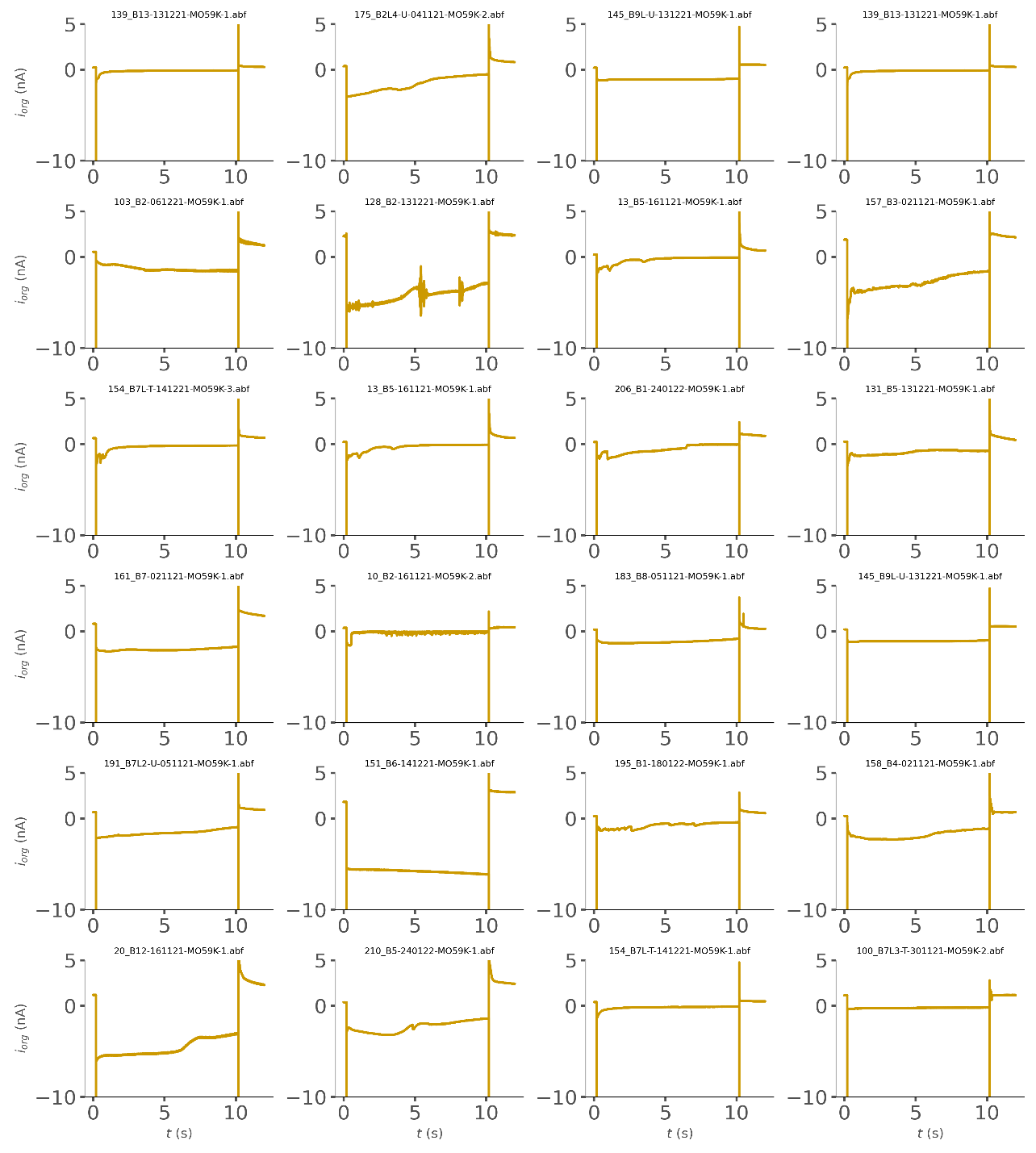

**Figure SF1.6.** Representative traces for the ion current i_org_ in the organic barrel when the potential is switched to V_org_=-500 mV for 10 s during the nanobiopsy phase.

### **S2: Longitudinal nanobiopsies of the same glioblastoma cells**

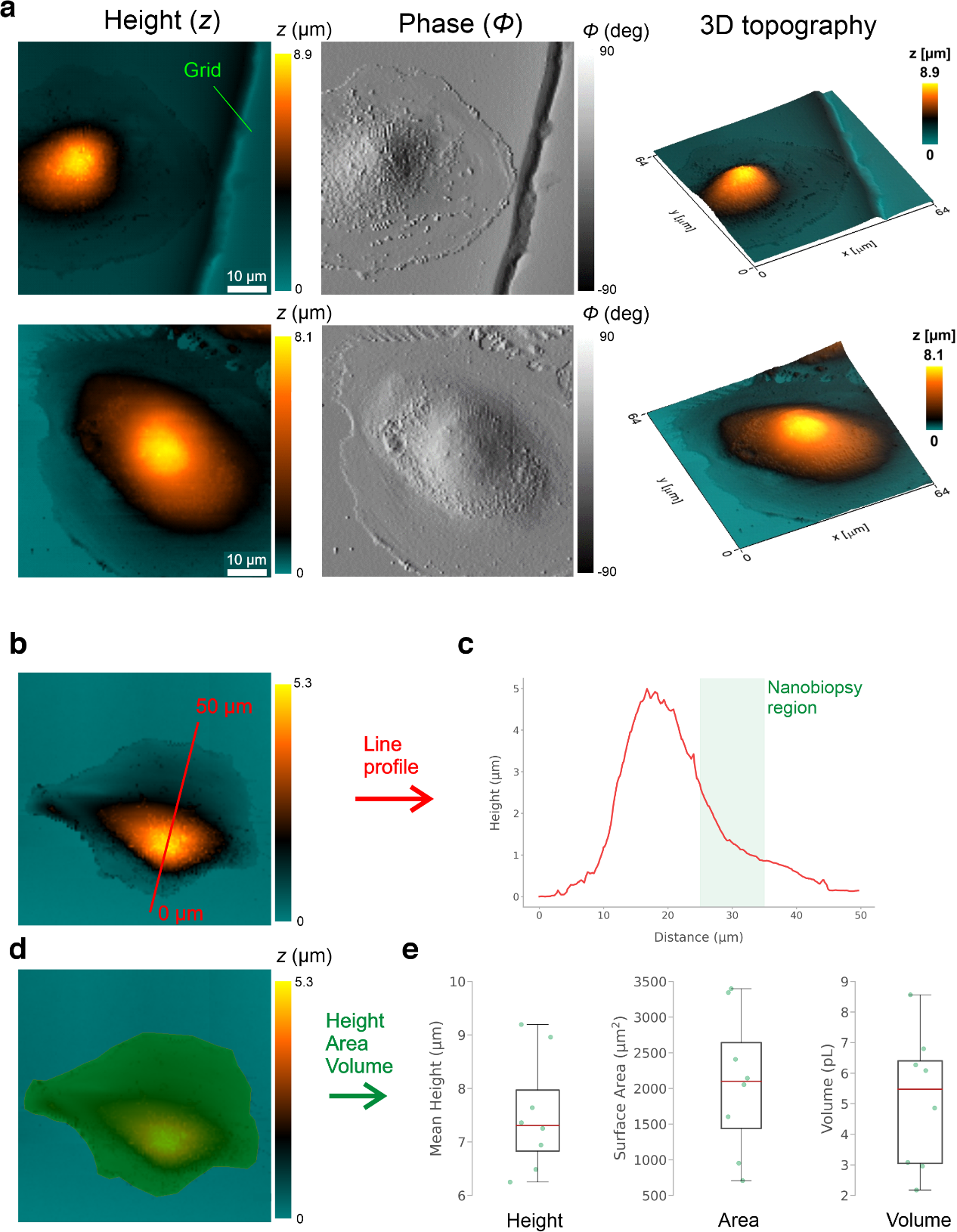

#### *S2.1 Determination of cell topography*

**Figure SF2.1.** Determination of cell topography and morphological parameters using SICM. (a) Surface topography measurements of live glioblastoma cells using SICM. Topographical maps were reconstructed using the height (left) and phase (center) channel. The height channel was used to generate a 3D image (right). The channel of the grid on the dish is visible in the topographical map of the first cell (top). (b) Topographical map of a single glioblastoma cell where a 50 µm line segment was traced (red) over the cytoplasm and nucleus to measure the cell height profile. (c) The high-slope region (green shaded area) was selected as suitable for nanobiopsy due to the slope facilitating the membrane penetration phase. A cell mask can be designed to select the pixels belonging to the cell (d) that are used to extract the mean height, surface area and volume whose values are shown in the boxplots (e) for a total number of cells equal to 8

#### *2.2 Fluorescent staining of fixed M059K GBM cells.*

Different combinations of dyes were used to image the organelles in the perinuclear region which was determined the most suitable region for the nanobiopsy. M059K cells were fixed using 4% paraformaldehyde and combinations of DAPI, Concanavalin A (Con A), MitoTracker green and cellMask orange were used to stain and image nuclear DNA, endoplasmic reticulum (ER), mitochondria and plasma membrane, respectively. **FIGURE SF2.2** shows the fluorescent micrographs obtained by combining DAPI, Con A, MitoTracker (a), DAPI, Con A (b) and DAPI, cellMask Orange (c).

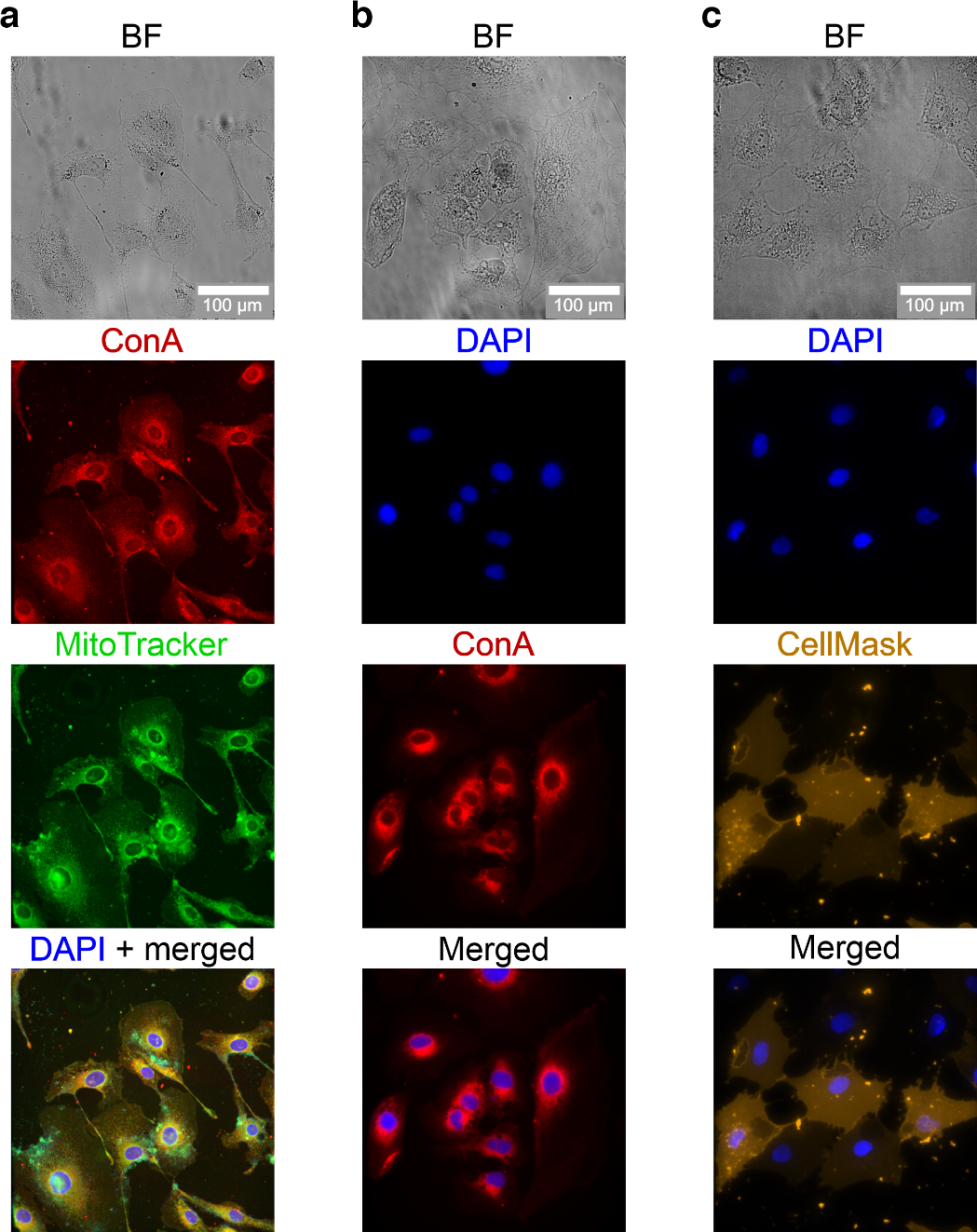

**Figure SF2.2:** Fluorescent staining of fixed M059K GBM cells. From top to bottom, (a) bright field BF, Con A, MitoTracker, merged ConA and MitoTracker and DAPI for nuclear DNA, endoplasmic reticulum and mitochondria visualisation. (b) BF, DAPI, ConA for nuclear DNA and endoplasmic reticulum visualisation. (c) BF, DAPI, cellMask orange for nuclear DNA and plasma membrane visualisation. Images were aquired using the BF, DAPI, TxRed and FITC filter sets.

#### *2.3 Experimental setup and longitudinal tracking of individual M059K GBM cells*

The experimental approach introduced in **FIGURE 4a,** **main text**, requires tracking the same cell over the time course of 72 hours. To assess whether a mixed culture of M059K_GFP_ and M059K_WT_ and the use of gridded dishes could be used for this purpose, we measured the cellular migration over 4 days. M059K_GFP_ and M059K_WT_ GBM cells were plated with a 1:350 ratio (M059K_GFP_ : M059K_WT_) according to the experimental methods in three different gridded dishes with the same seeding density ($2.5\cdot{10}^{4}$ cells/dish) on day 0 of the experiment. 24 hours later (day 1), an individual M059K_GFP_ was identified and imaged in a specific location of the grid, ensuring that only one M059K_GFP_ was present in the field of view imaged with a 10x magnification objective. After imaging, the dish was brought back to the incubator and the same location was imaged again after 24 (day 2), 48 (day 3) and 72 hours (day 4). The gridded dish enabled tracking of the same cell over time by finding back the same labelled position on the grid where the M059K_GFP_ cell was identified on day 1. **FIGURE SF2.3a** shows one example of an individual M059K_GFP_ cell imaged on day 1 in position A11 on the grid. After 24 hours (day 2), the same cell was found back by imaging the same dish location (A11). Two cells were found on day 2, suggesting that the cell went through cell division. After 48 (day 3) and 72 hours (day 4) the same cells were found back in the same location. On day 4, four cells were found around A11, suggesting that both cells found on day 2 and day 3 divided. An image processing routine was developed to measure different cell migration parameters:

- The **total cell migration Δ*x*_tot_** is the maximum distance travelled by the cell over 72 hours and it was calculated as the sum of the longest distance (even if the least likely) travelled by the cell every 24 hours (**Δ*x*_1,_ Δ*x*_2_**, **Δ*x*_3_),** as shown in **FIGURE 2.3b**
- The **maximum cell migration Δ*x*_max_** was calculated as the length of the segment (orange) connecting the point indicating the cell on day 1 (purple) to the farthest point indicating the farthest cell on day 4 (red), as shown in **FIGURE SF2.3c**
- The **total area enclosing all cells after 72 hours *A*_all_** was measured by tracing the smallest rectangle enclosing all the points/cells of all days (purple, green, blue, red), as shown in **FIGURE SF2.3d**
- The **mean migration over 24 hours** was measured by considering all segments connecting points/cells across 24 hours and calculating the mean value.
- The **field of view (FOV)** is defined as the observable area under the microscope using a 10x objective. The field of view shown in **FIGURE SF2.3e** (black) has size 1300 x 1300 µm and area 1.69 mm^2^.

The parameters listed above were measured for three cells across three different dishes. **FIGURE SF2.3f-i** shows the results obtained for each of these parameters. For all parameters, no statistical significance was found between different replicates (t-test, p<0.05) except from the total migration between dish 1 and dish 2 which showed a slight significance (p=0.05). The total migration Δ*x*_tot_ (**FIGURE SF2.3f**) and the maximum migration Δ*x*_max_ (**FIGURE SF2.3g**) show that the longest distance that a cell can travel in the worst-case scenario is always significantly smaller than the dimensions of the field of view. This suggests that a cell is not able to migrate out of the field of view in the time course of the experiment. The enclosing area *A*_all_ (**FIGURE SF2.3h**) was found in all cases significantly smaller than the area of the field of view. The biggest *A*_all_ found had value 0.35 mm^2^ which is ~5 time smaller than the area of the field of view (1.69 mm^2^), suggesting that all cells (initial cell and progeny) remain within the field of view area in the time course of the experiment. The mean migration over 24 hours (**FIGURE SF2.3i**) also suggests that daily distance travelled by the cell is not enough to result into the cell exiting the field of view.

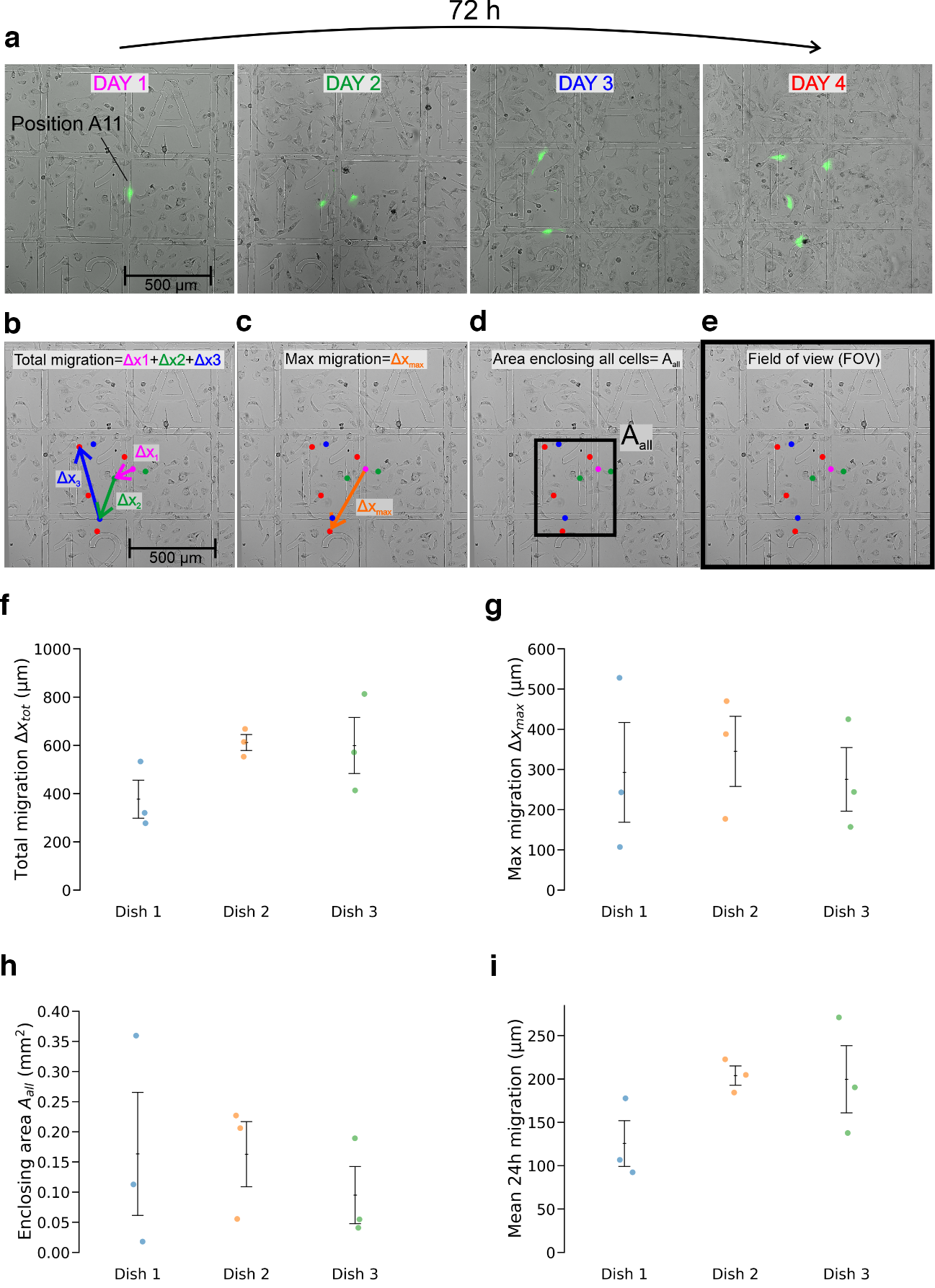

**Figure SF2.3.** Analysis of cell migration and division over 72 hours. (a) a single GFP-transfected cell is identified on day 1 in a specific location on the grid (A11) and it is tracked over 72 hours (DAY1-DAY4). An image processing routine allowed the quantification of the total cell migration (b), the maximum cell migration over 72 hours (c), and the area enclosing all cells after 72 hours (d). The field of view (FOV) defines the observable area, and it is imposed by the microscope objective. The biggest field of view available for this study was 1300 x 1300 μm. The total cell migration (a), the maximum migration over 72 hours (b), the area enclosing all cells after 72 hours (c), and the mean cell migration over 24 hours (d) were measured from 3 cells across 3 different dishes. All parameters suggest that the extent of cell migration does not exceed the dimensions of the field of view.

#### *2.4 Longitudinal sampling of the same glioblastoma cell*

To determine whether the same cell could be sampled at different time points, a mixed culture of M059K_GFP_ and M059K_WT_ was plated on a gridded dish, as described in section SF2.3, and an individual M059K_GFP_ was identified near the position H20 on the grid using a 10x magnification objective, ensuring the presence of only one M059K_GFP_ in the field of view. Next, a nanobiopsy was collected resulting in the cell emitting a red fluorescence signal (false yellow colour) following nanoinjection of the red fluorophore ATTO 565 (**FIGURE SF2.4a**). The dish containing the nanobiopsied cell was brought back to the incubator and the cell was imaged again after 24 hours (day 2, **FIGURE SF2.4b**) and 48 hours (day 3, **FIGURE SF2.4c**) and was found approximately in the same position (H20) on the grid. After 72 hours (day 4), a longitudinal nanobiopsy was collected resulting in the cell emitting a red fluorescence signal following nanoinjection (**FIGURE SF2.4d**).

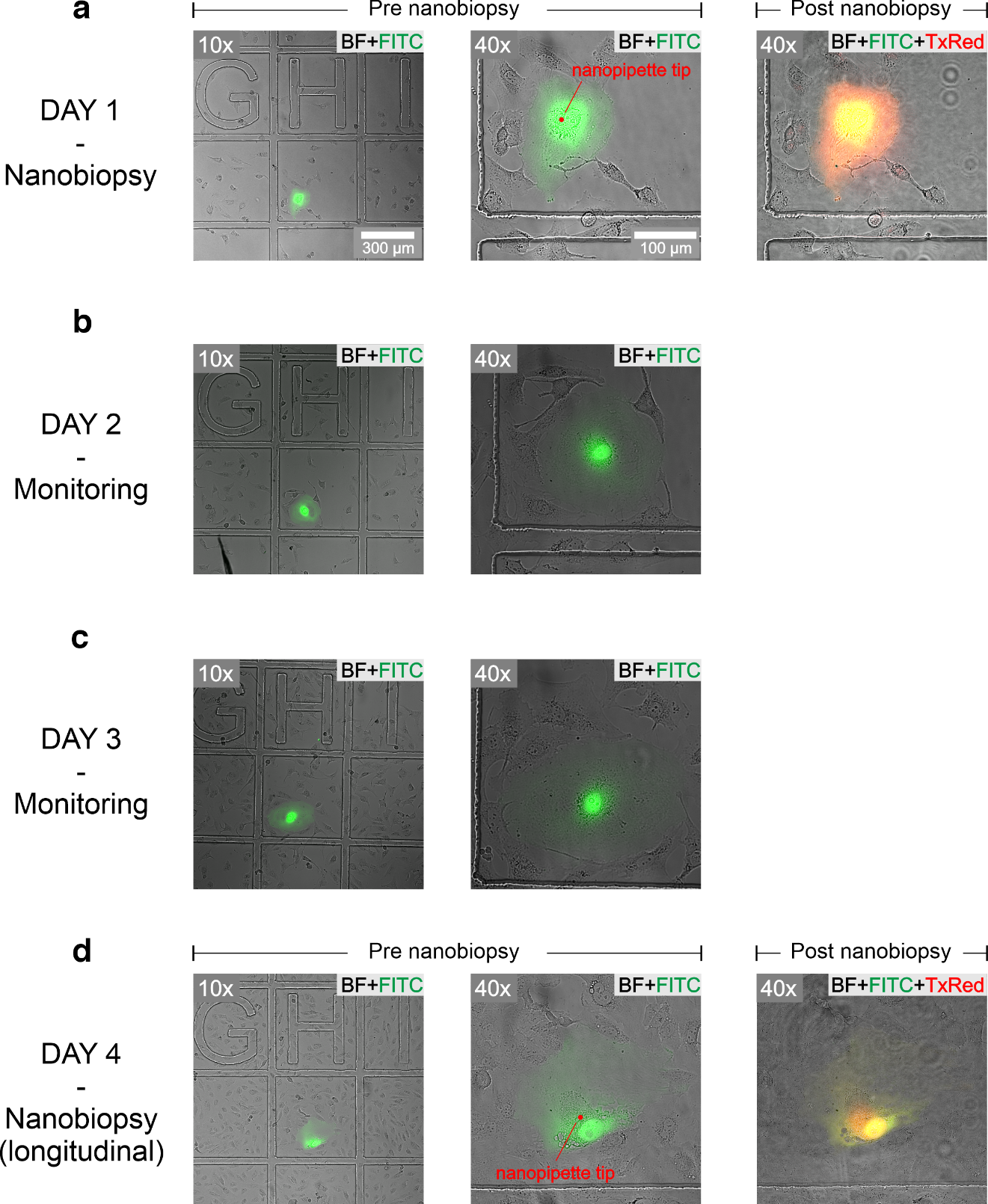

**Figure SF2.4.** Longitudinal nanobiopsy of the same cell. On day 1 (b), a nanobiopsy was collected from an individual M059K_GFP_ localised in position H20 on the grid resulting in the cell emitting a red fluorescent signal following nanoinjection of a red fluorophore (false yellow colour). The cell was imaged again on day 2 (b) and day 3 (c) when it was found around the same position H20. On day 4 (d), a longitudinal nanobiopsy was performed resulting in the cell emitting a red-fluorescent signal following nanoinjection. Micrographs were obtained using the filter set for bright field (BF), fluorescein isothiocyanate (FITC) and texas red (TxRed). The fluorophore injected was ATTO 565.

#### *2.5 Examples of longitudinal nanobiopsies on untreated cells*

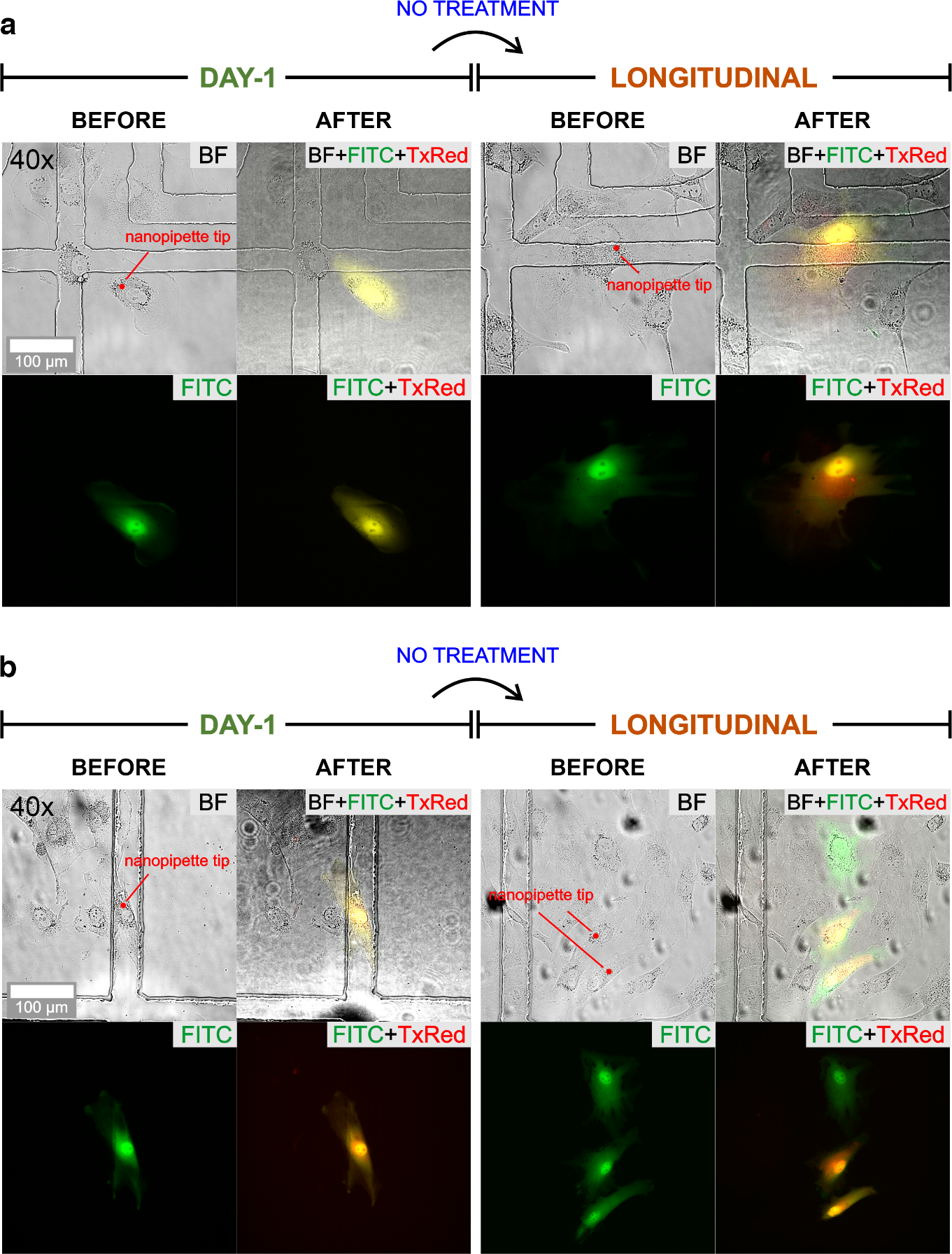

**Figure SF2.5.** Examples of longitudinal nanobiopsies collected from untreated M059K GBM cells. (a) Optical (BF) and fluorescence (FITC, TxRed) micrographs of an individual M059K_GFP_ cell that is nanobiopsied and nanoinjected on day 1 that is left untreated and nanobiopsied and nanoinjected again on day 4 (longitudinal). (b) Optical and fluorescence micrographs of an individual M059K_GFP_ cell that is nanobiopsied and nanoinjected on day 1, survive over 72 hours and divide, whose progeny is nanobiospied and nanoinjected a second time on day 4.

#### *2.6 Quality control of multiplexed single-cell libraries*

Two multiplexed libraries were generated for the whole-cell lysate (positive control) and nanobiopsy samples, respectively. A tape station was used to determine the fragment size of both libraries. In both cases, the fragment size was within the range recommended by the manufacturer.

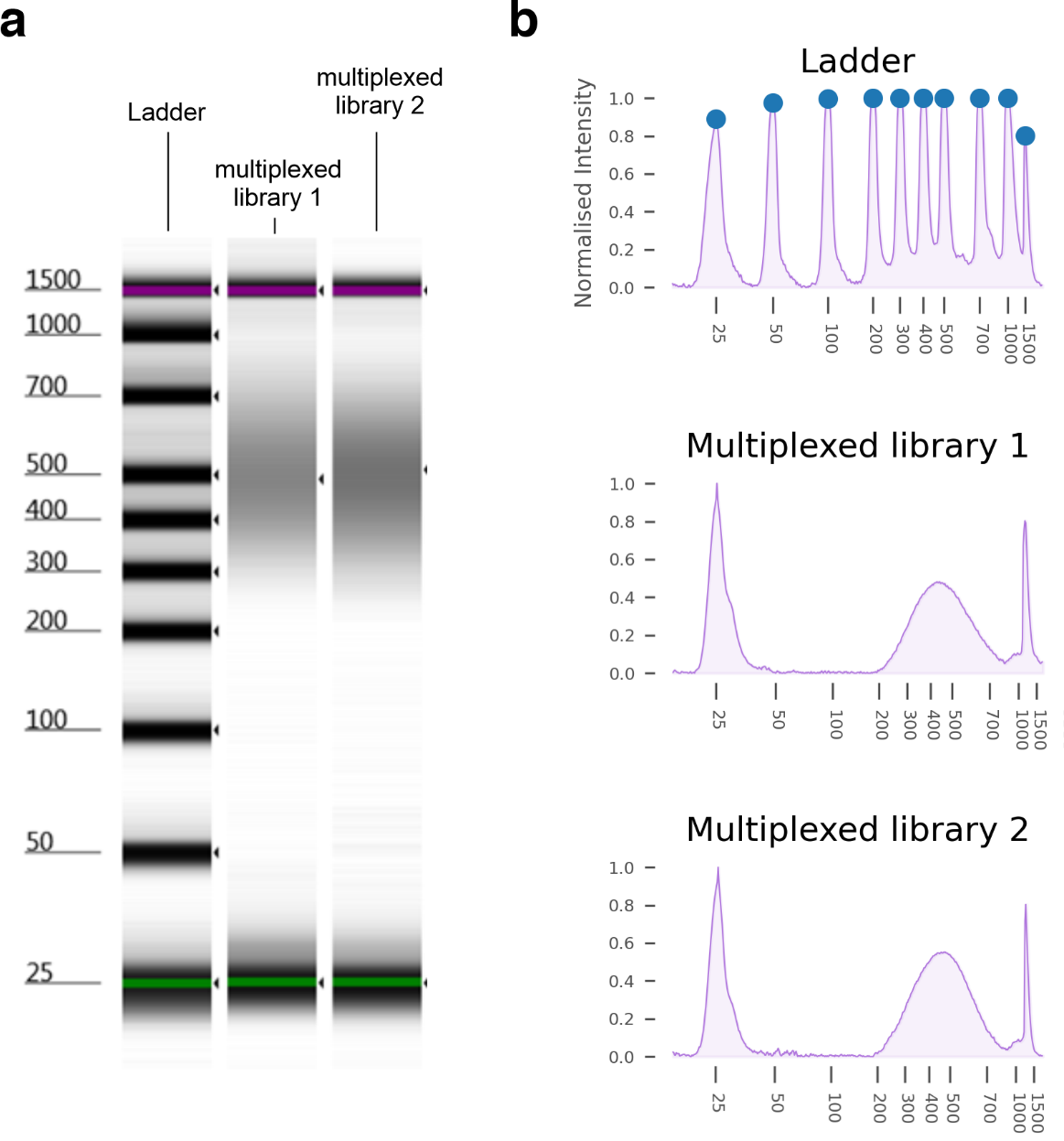

**Figure SF2.6.** Quality control of multiplexed single-cell libraries. (a) Tape station traces showing the calibration ladder, the multiplexed library 1 containing the whole-cell lysate samples and the multiplexed library 2 containing all the nanobiopsy samples. (b) densitometry traces extracted from the tape station images showing the fragment size distribution of both multiplexed libraries.

### **S3: Sequencing data analysis**

#### *3.1 Quality metrics of unfiltered data*

**Figure SF3.1:** Main sequencing metrics before data filtering of day-1 nanobiopsy (NB_Day1) longitudinal nanobiopsy of treated (NB_T) and untreated (NB_U), and whole-cell lysate treated (WC_T) and untreated (WC_U) samples.

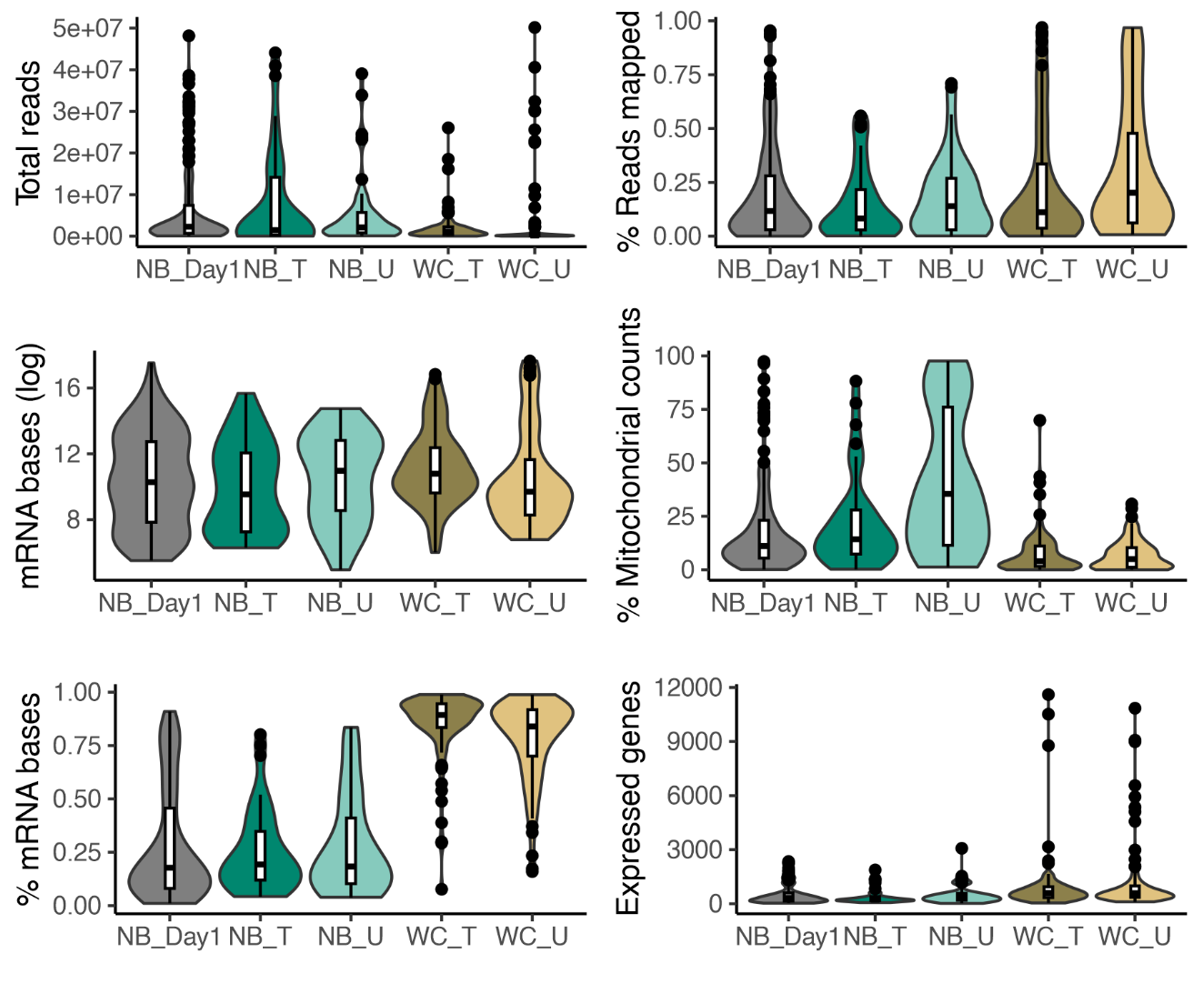

#### *3.2 Principal component analysis visualisation of longitudinal nanobiopsy data.*

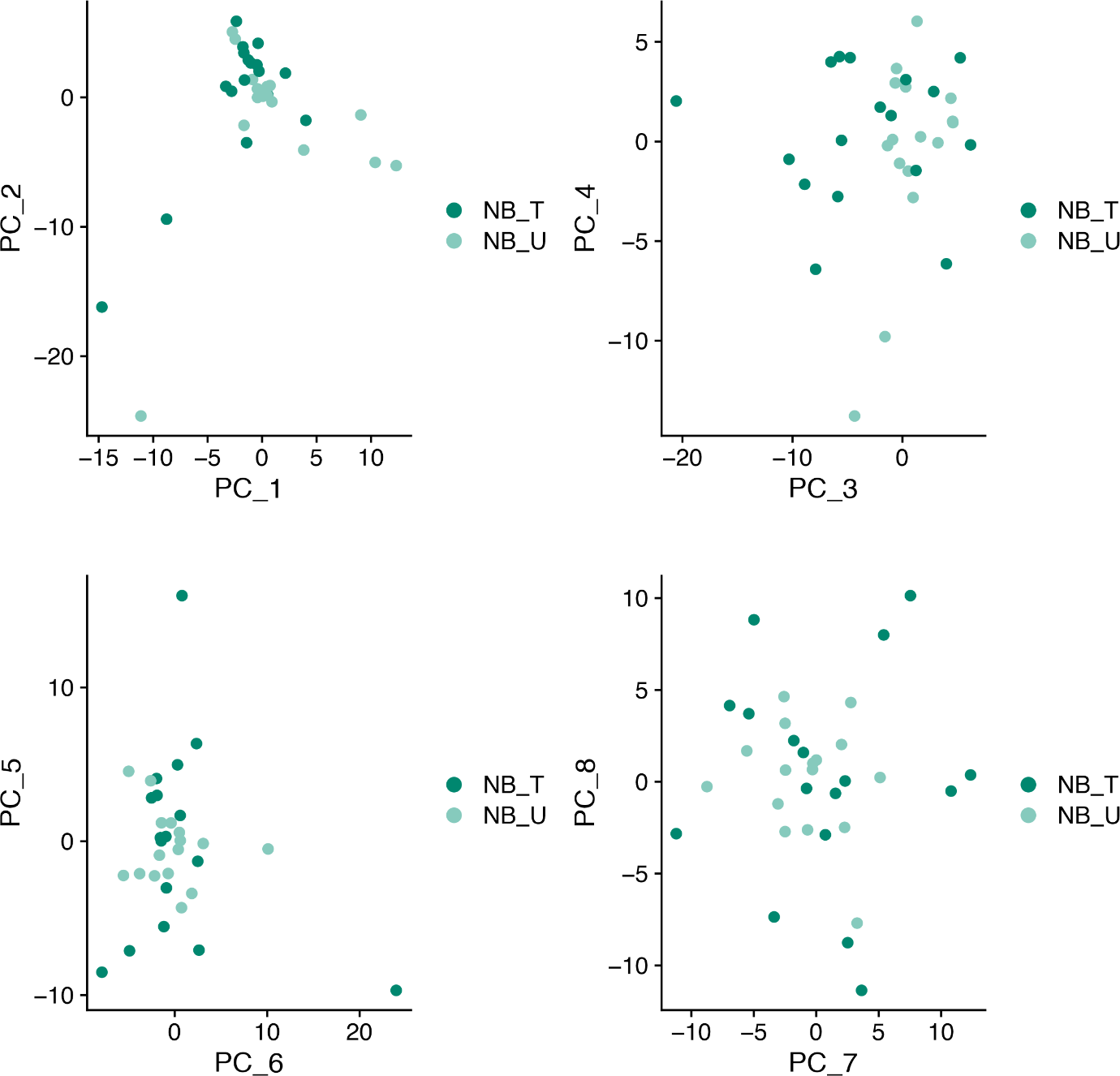

**Figure SF3.2:** Visualisation of longitudinal nanobiopsy data of treated (NB_T) and untreated (NB_U) samples along the first eight principal components.

#### *3.3 Visualisation of lack of bias introduced by the metrics on the UMAP plot.*

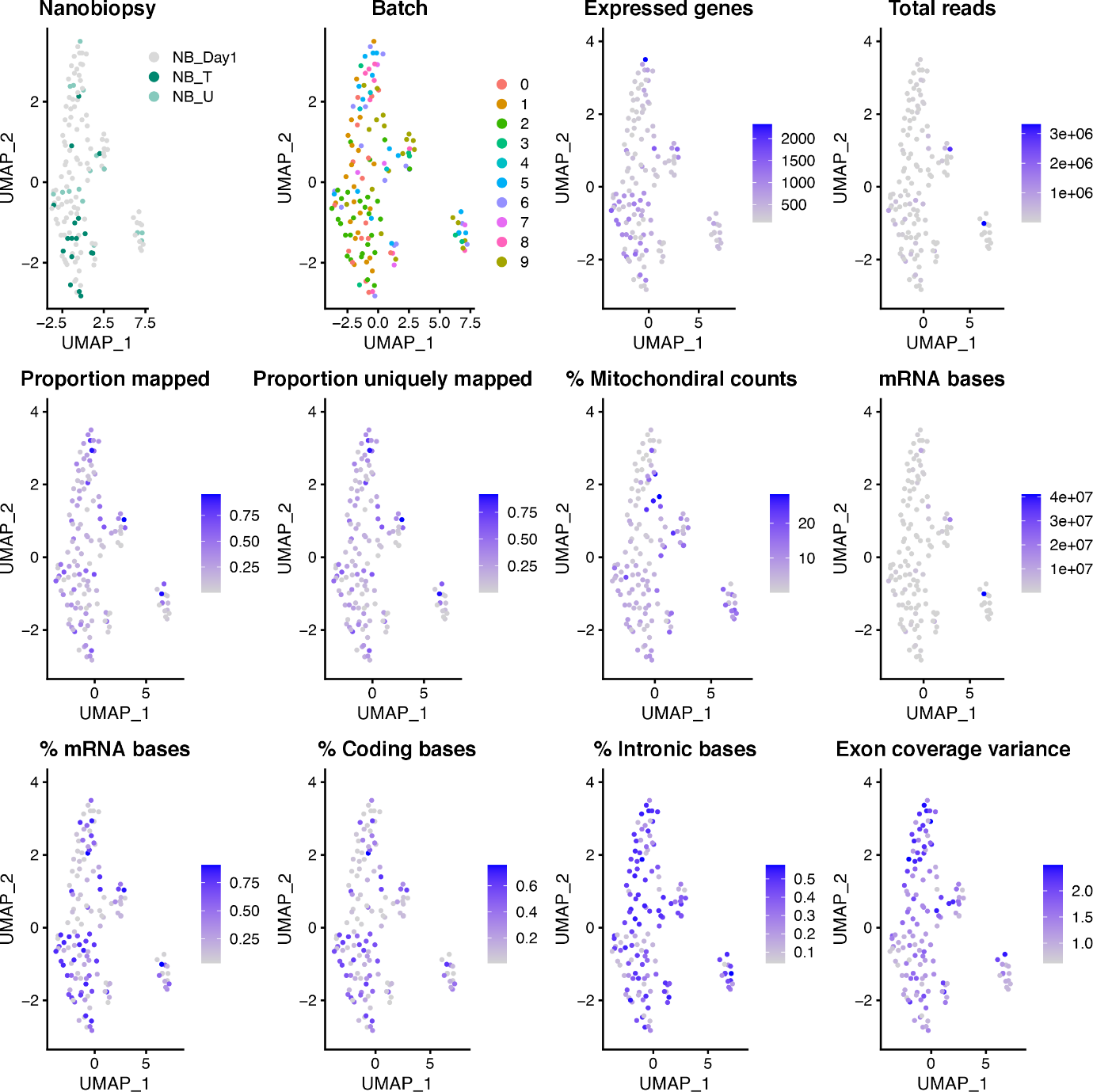

**Figure SF3.3:** Visualisation of technical bias due to batch and sequencing metrics in the UMAP plot.

#### *3.4 Nanobiopsy count after filtering and GSVA scoring*

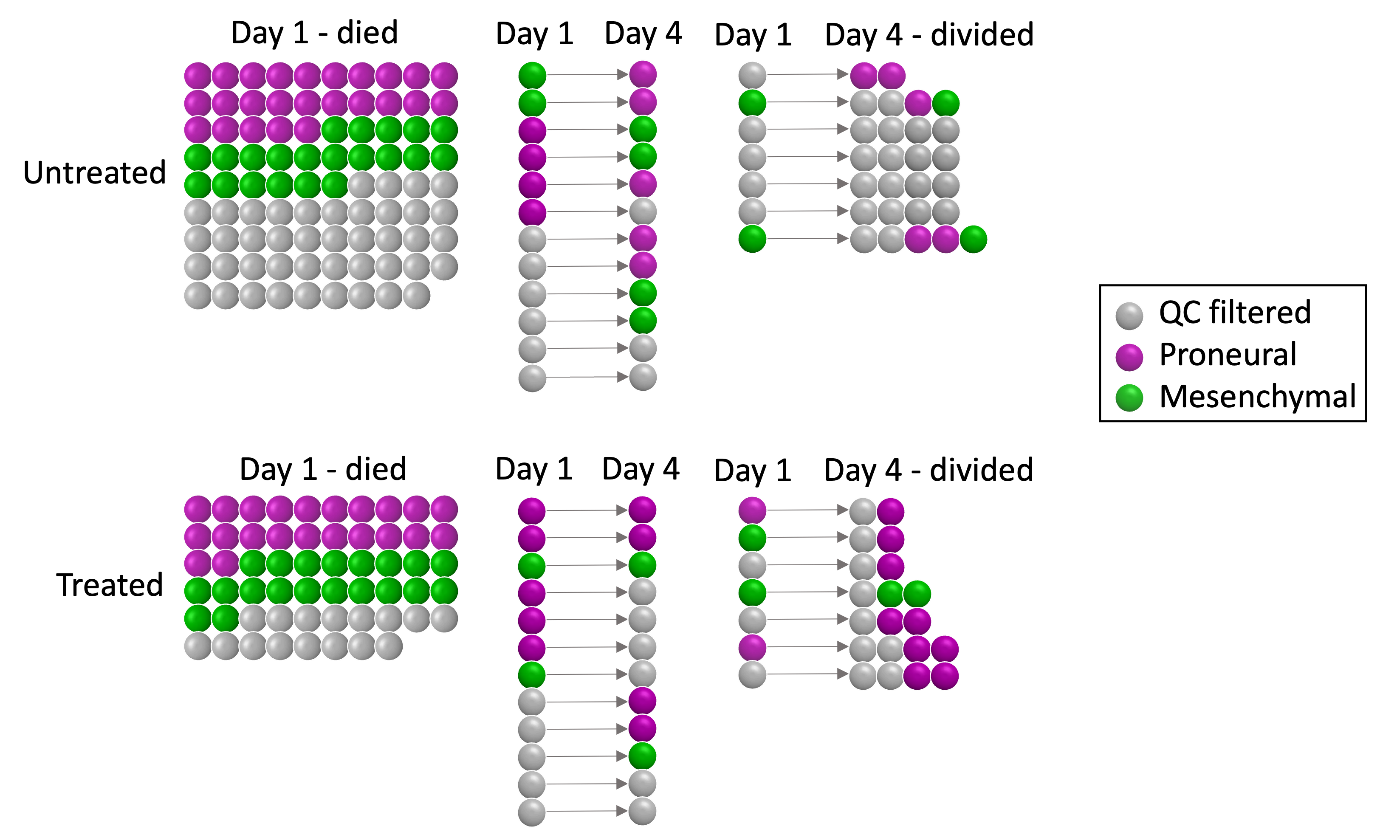

**Figure SF3.4:** Nanobiopsy count after filtering and GSVA scoring. 55% of the nanobiopsy samples met the filtering criteria and were classified in the mesenchymal (green) and proneural (purple) GBM subtype.

#### *3.5 Change in the proneural and mesenchymal score of treated and untreated longitudinal nanobiopsy samples.*

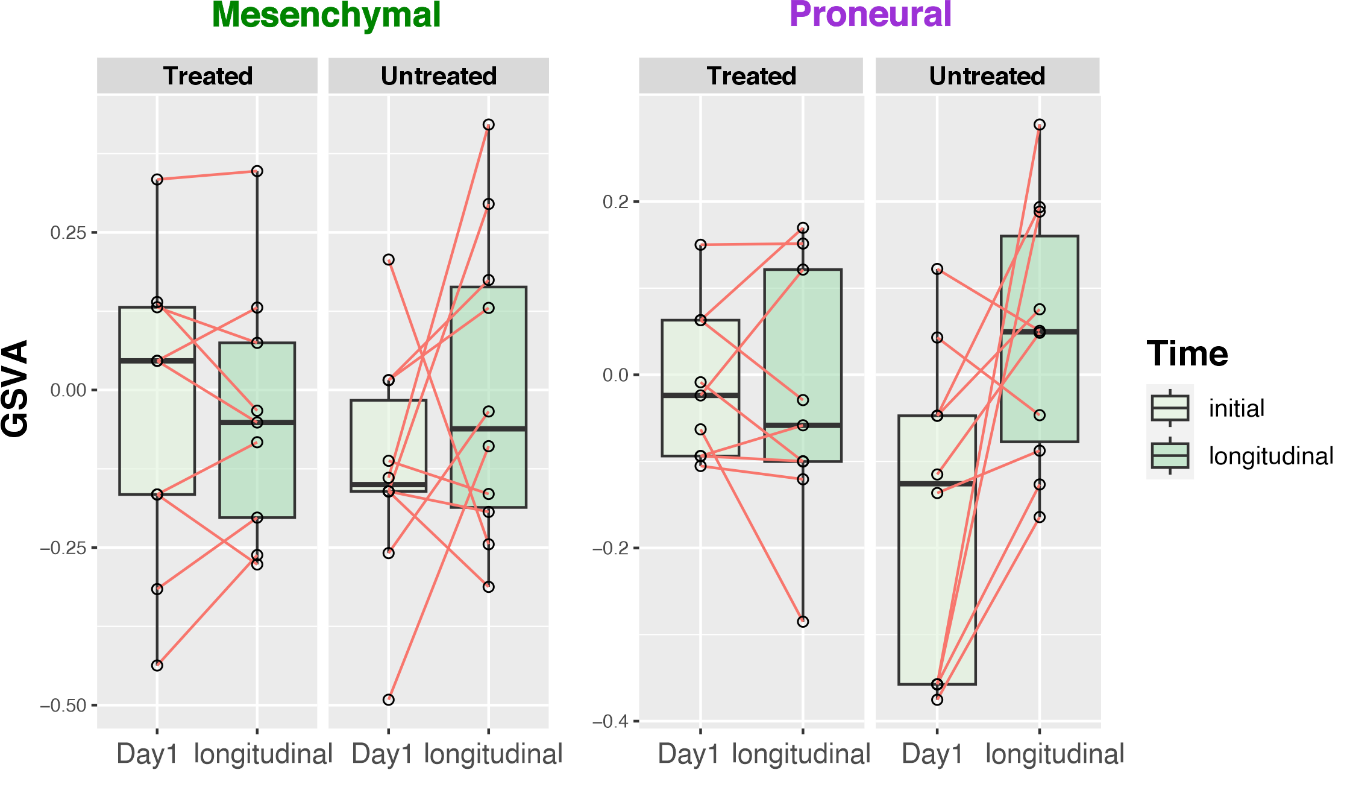

**Figure SF3.5.** Boxplots showing the individual GSVA score for the mesenchymal (green) and proneural (purple) subtype of paired day1 and longitudinal treated and untreated nanobiopsy samples.
